# Ras-dependent activation of BMAL2 regulates hypoxic metabolism in pancreatic cancer

**DOI:** 10.1101/2023.03.19.533333

**Authors:** Alvaro Curiel-Garcia, Sam R. Holmstrom, Cristina Castillo, Carmine F. Palermo, Steven A. Sastra, Anthony Andren, Lorenzo Tomassoni, Li Zhang, Tessa Y.S. Le Large, Irina Sagalovskiy, Daniel R. Ross, Winston Wong, Kaitlin Shaw, Jeanine Genkinger, Hanina Hibshoosh, Gulam A. Manji, Alina C. Iuga, Roland M. Schmid, Kristen Johnson, Michael A. Badgley, Pasquale Laise, Costas A. Lyssiotis, Yatrik M. Shah, Andrea Califano, H. Carlo Maurer, Kenneth P. Olive

## Abstract

KRAS is the archetypal oncogenic driver of pancreatic cancer. To identify new modulators of KRAS activity in human pancreatic ductal adenocarcinoma (PDAC), we performed regulatory network analysis on a large collection of expression profiles from laser capture microdissected samples of PDAC and benign controls. We discovered that BMAL2, a member of the PAS family of transcription factors, promotes tumor initiation, progression, and post-resection survival, and is highly correlated with KRAS activity. Functional analysis of BMAL2 target genes suggested a role in regulating the hypoxia response, a hallmark of PDAC. Knockout of *BMAL2* in multiple human PDAC cell lines reduced cancer cell viability, invasion, and glycolysis, leading to broad dysregulation of cellular metabolism, particularly under hypoxic conditions. We find that BMAL2 directly regulates hypoxia-responsive target genes and is necessary for the stabilization of HIF1A under low oxygen conditions, while simultaneously destabilizing HIF2A. Notably, *in vivo* xenograft studies demonstrated that BMAL2 loss significantly impairs tumor growth and reduces tumor volume, underscoring its functional importance in tumor progression. We conclude that BMAL2 is a master transcriptional regulator of hypoxia responses in PDAC that works downstream of KRAS signaling, possibly serving as a long-sought molecular switch that distinguishes HIF1A- and HIF2A-dependent modes of hypoxic metabolism.

**Statement of Significance:** We annotate the landscape of KRAS-associated transcriptional drivers of pancreatic cancer initiation, progression, and overall survival, leading to the identification of BMAL2 as a novel regulator of hypoxic metabolism. BMAL2 helps execute the oncogenic transcriptional programs of KRAS and serves as a long-sought switch between HIF1A- and HIF2A-dependent modes of hypoxic metabolism.

## Introduction

DNA sequencing of hundreds of human pancreatic tumors has helped define the genetic drivers of pancreatic ductal adenocarcinoma (PDAC). However, mutations alone poorly predict key, clinically-relevant traits of the disease, such as tumor stage or therapeutic response ^1,2^. This suggests that non-genetic factors may control vital, biological characteristics of PDAC, such as differentiation state, metastasis, and clinical outcome. Defining these factors is a necessary first step towards intervening in these complex pathologies.

Among genetic drivers, activating mutations in *KRAS* are the most penetrant, driving ∼95% of human PDAC tumors. Though extensive work has delineated the signal transduction pathways that effect its activity, there is a comparatively poor understanding of how the downstream transcriptional outputs of mutant *KRAS* drive contribute to the manifold phenotypes attributed to RAS activity. While MYC and AP1 serve as canonical effectors of RAS signaling, hyperactivation of these transcription factors alone does not recapitulate the tumorigenic phenotype of *Kras* mutation in the pancreas ^3,4^. The advent of RAS inhibitors for clinical use has served to highlight the need for a more detailed understanding of how RAS signaling drives pancreatic tumors, not only from a classical genetic standpoint, but from a system-wide, cellular view.

RNA sequencing (RNA-Seq) has been widely used to identify correlations between gene expression and phenotypes. However, recent advances in the area of regulatory network analysis have enabled the identification of proteins that causally drive phenotypes using RNA expression data ^5^. Briefly, this approach quantifies the signaling activity of transcription factors and other transcriptional regulators (collectively, “regulatory proteins” or RPs) by integrating the expression of their positive and negative target genes using algorithms such as VIPER ^6,7^. This approach is founded on algorithms such as ARACNe, which can accurately infer context-specific sets of target genes for thousands of RPs using context-specific gene expression datasets ^8–10^. Within this framework, master regulators (MRs) are a distinct subset of RP whose activity is both necessary and sufficient to drive specific cellular phenotypes. Here we apply regulatory network analysis to a set of 242 laser capture microdissected samples of human PDAC or precursor lesions in order to understand how mutant *KRAS* drives malignant phenotypes in PDAC on a comprehensive scale.

## Results

### BMAL2 Drives PDAC Phenotypes

To link clinical and pathological phenotypes to gene expression data in PDAC, we first augmented our previous collection of RNA-Seq profiles derived from the malignant epithelium of laser-captured, microdissected (LCM) human PDAC (CUMC-E, ^11^) to now include a total of 197 adenocarcinomas, 26 low-grade pancreatic intraepithelial neoplasms (PanINs), and 19 low-grade intraductal papillary mucinous neoplasms (IPMNs) ^12^. Together, the PanIN and IPMN samples served as “benign controls” that are committed to the neoplastic lineage ^13,14^ but unlikely to progress to PDAC ^12,15^. Each sample was associated with clinical data including demographics, surgical features, treatment class, survival time, and histopathological analysis performed on a section adjacent that used for LCM. Unsupervised clustering by Principal Component Analysis (PCA) showed a clear distinction between precursor and PDAC expression profiles (**Figure 1A**).

**Figure 1.**
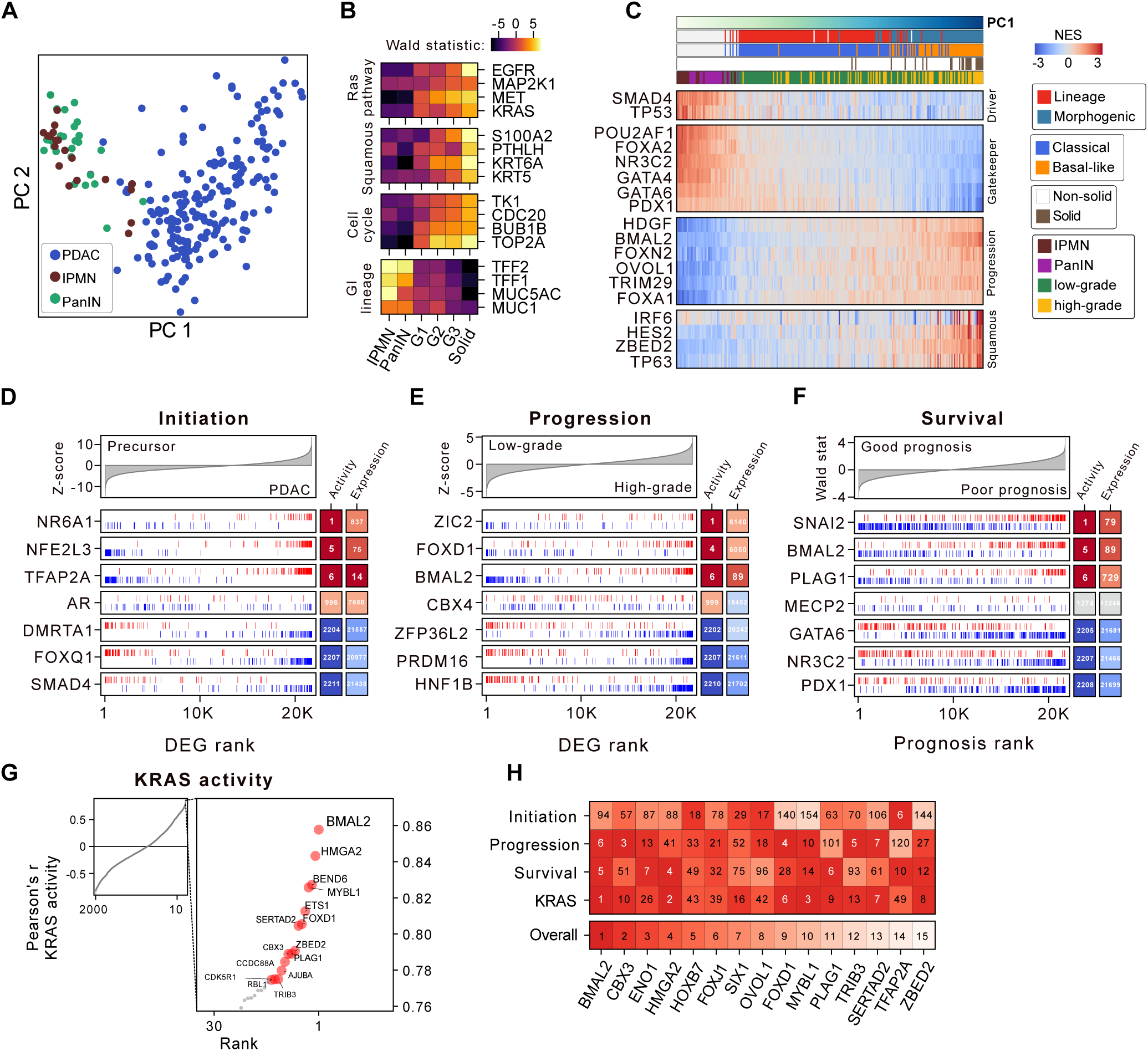
Master regulator analysis of PDAC nominates BMAL2 as a PDAC driver **(A)** Principal component analysis (PCA) of precursor and PDAC LCM RNA-Seq samples used to assemble a regulatory model (interactome) of PDAC carcinogenesis. (**B**) Heatmap depicting select differentially expressed genes between the indicated histopathological groups (x-axis). Wald test statistics were derived from a negative-binomial linear model comparing the respective group against all other groups. (**C**) Heatmap of protein activity scores (NES) for select regulatory proteins during PDAC progression. The samples are ordered by their value in the first principal component, essentially capturing progression and dedifferentiation. (**D**) Select results from master regulator analysis on a genome-wide PDAC initiation gene expression signature (x-axis) represented by Z-scores for each gene. Each regulatory protein’s regulon is represented by red (positive targets) and blue (negative targets) vertical bars. The rank of each RP based on activity and expression, respectively, is illustrated on the right. (**E**) Select results from master regulator analysis on a genome-wide PDAC progression gene expression signature (x-axis) (**F**) Select results from master regulator analysis on a genome-wide survival signature (x-axis) represented by Wald test statistics from a multivariate Cox Proportional Hazards model testing the coefficient for each gene’s continuous expression while accounting for patient age (**G**) Genome-wide Pearson correlation between KRAS and RP activity with illustration of the most positively correlated RPs (**H**) Heatmap of rank positions for the indicated regulatory proteins (x-axis) in each of four critical phenotypic PDAC transitions yields a conserved core set of non-oncogene dependency candidates for PDAC. A low rank represents activation in a given phenotype signature.

To benchmark the dataset, we conducted an unbiased analysis of associations between gene expression and histopathological features, as drawn from observations of adjacent tissue sections (**Supplementary Figure S1A**). We found that poorly differentiated tumors had elevated expression of *KRAS* ^16,17^, of proliferation markers (*TOP2A*, *BUB1B*, *CDC20*, and *TK1*) ^18^, and indicators of squamous PDAC lineage (*KRT5*, *KRT6A*, *PTHLH*, and *S100A2*) ^19,20^ (**Figure 1B**). Conversely, hallmarks of differentiation (*MUC1*, *MUC5AC*) and GI lineage (*TFF1*, *TFF2*) decreased during tumor initiation and progression, providing strong validation from these established biomarkers.

While correlations identified through differential gene expression analysis can yield some insights, regulatory network analysis can connect phenotypes to their mechanistic drivers ^6,8,10,21,22^. We therefore applied ARACNe ^6,7^ to the full set of 242 epithelial expression profiles to generate a regulatory network specific to PDAC epithelia, compiling a total of 263,085 inferred transcriptional targets for 2,211 regulatory proteins (RP). For 26 of these RPs, ChIP-Seq data were publicly available in human PDAC cells ^23^. Half of these (13/26) showed a significant overlap with the ARACNe target gene sets, despite the difference of *in vivo* versus *in vitro* contexts (**Supplementary Figure S1B**), providing experimental validation for the accuracy of the regulatory network. Finally, to benchmark the regulatory network against established PDAC biology, we examined the inferred activities of RPs with well-studied roles in PDAC (**Figure 1C**). Principal component 1 (PC1) effectively captured the progression from benign precursors through low-grade PDAC, to high-grade PDAC with squamous features. These phenotypic changes were accompanied by the repression of canonical PDAC tumor suppressors TP53 and SMAD4, and activation of RPs with known oncogenic functions in PDAC, such as FOXA1 ^24^ and TRIM29 ^25^ (**Figure 1C**). GI transcription factors GATA6, FOXA2, and PDX1, which have been described as being “overexpressed” in some PDAC molecular subtypes ^17,26^, were more highly active in low-grade PDAC than in high grade. However, in comparison to benign precursors, their activity in low-grade tumors was down-regulated, consistent with the progressive loss of GI identity during tumor initiation and progression. Finally, drivers of squamous histology such as TP63 ^19^ and ZBED2 ^27^ were hyperactivated in high-grade PDAC, particularly those with annotated squamous histopathology. Together these findings demonstrate the ability of MR analysis to accurately identify known drivers of specific PDAC phenotypes.

Next we examined the RPs whose activities were most associated either with KRAS activity or with key malignant phenotypes, including tumor initiation, tumor progression, and patient survival (**Supplementary Figure 1C)**. We used MARINa analysis ^28^ to identify MRs of PDAC initiation (comparing precursors to adenocarcinoma, **Figure 1D**) and progression (comparing PDAC with low-grade versus high-grade histopathology, **Figure 1E**). For survival, we constructed a survival signature from a multi-variate Cox proportional hazards model and identified RPs controlling the expression of the most prognostic target genes (**Figure 1F**). Lastly, we calculated KRAS activity by iterating a new PDAC regulatory network comprising the transcriptional targets for a total of 2,523 signaling factors (see Methods) (**Figure 1G**). In this analysis, the inferred target genes of each signaling protein serve as a bespoke reporter gene set, providing an indirect, but unbiased, measure of their signaling activity in PDAC ^9,29^. As expected, the inferred activity of KRAS increased significantly during tumor initiation and tumor progression (**Supplementary Figure 1D**), peaking among tumors growing in solid nests.

The positive and negative MRs of all four of these phenotypes were widely associated with the oncogenic programs of the HALLMARK signature set ^30^ **(Supplementary Figure S1E)**. However, in contradistinction to the processes of PDAC initiation and progression, overall survival was not associated with proliferation gene sets. Rather, MRs of patient survival were most strongly associated with hypoxia, KRAS signaling, immune signaling, and EMT, suggesting that clinical outcome is not solely dependent on growth rate ^31^. To identify new global drivers of PDAC malignancy, we integrated the ranked lists of MRs for PDAC initiation, progression, survival, and KRAS activity. We found that BMAL2, a member of the PAS superfamily, was the top candidate master transcriptional regulator of PDAC malignancy (**Figure 1H**).

### BMAL2 is associated with aggressiveness in multiple PDAC datasets

BMAL2 has not previously been defined as a key driver of PDAC. To assess the reproducibility of our findings, we performed a meta-analysis across a total of 10 published PDAC expression studies ^26,32–40^ and found that *BMAL2* expression was elevated relative to normal pancreas (**Supplementary Figure S2A**) and elevated in high-grade versus low-grade PDAC specimens (**Supplementary Figure S2B**). Concordant with our findings in the CUMC-E cohort, high BMAL2 activity consistently identified patients with worse outcomes (**Supplementary Figure S2C**). Next, we evaluated each combination of five subtype classification schemes and six PDAC expression data sets for differences in BMAL2 activity between tumors of the most aggressive versus least aggressive subtype (**Supplementary Figure S2D**), and found that BMAL2 was consistently hyperactivated in the most aggressive subtype ^41–43^.

Finally, we found *BMAL2* expression in normal tissues (**Supplementary Figure S2E**) ^41^ to be highest in squamous epithelia, whereas expression in the normal pancreas was comparatively low. By contrast, PDAC tumor samples had among the highest levels of *BMAL2* expression across multiple cancers (**Supplementary Fig. S2F**) ^42^ and PDAC cell lines had the highest median expression across cell lines from different lineages ^43^ (**Supplementary Figure S2G**). Together these results validate our identification of BMAL2 as a key driver of initiation, progression, and outcome in multiple independent PDAC datasets.

### Oncogenic KRAS activates BMAL2 through ERK

In addition to driving the three PDAC malignancy phenotypes, BMAL2 stood out as the single RP most highly correlated with KRAS activity (out of 2211 measured), leading us to hypothesize that BMAL2 is regulated by KRAS signaling. To test this, we reanalyzed published expression datasets in which mutant KRAS activity was experimental manipulated to assess BMAL2 activity:

*Kras^G12D^*activation in murine pancreatic ductal cells (Sivakumar, Diersch); *Kras^G12D^* reactivation in PDAC cells that survived mutant *Kras* withdrawal (Viale); and mutant *Kras* inactivation in murine and human PDAC cells (Ying, Bryant) ^44–48^. In all five independent experiments, BMAL2 activity was regulated as predicted by KRAS, suggesting that BMAL2 may serve as a transcriptional effector downstream of mutant KRAS (**Figure 2A**). To determine whether this association extended beyond PDAC, we examined similar RAS modulation experiments in five datasets from lung adenocarcinoma models (LUAD) and three datasets from colorectal adenocarcinoma models (COAD). In all save one LUAD experiment, BMAL2 activity was significantly regulated in coordination with RAS (**Figure 2A**), suggesting a more general association of BMAL2 function in RAS-driven cancers.

**Figure 2.**
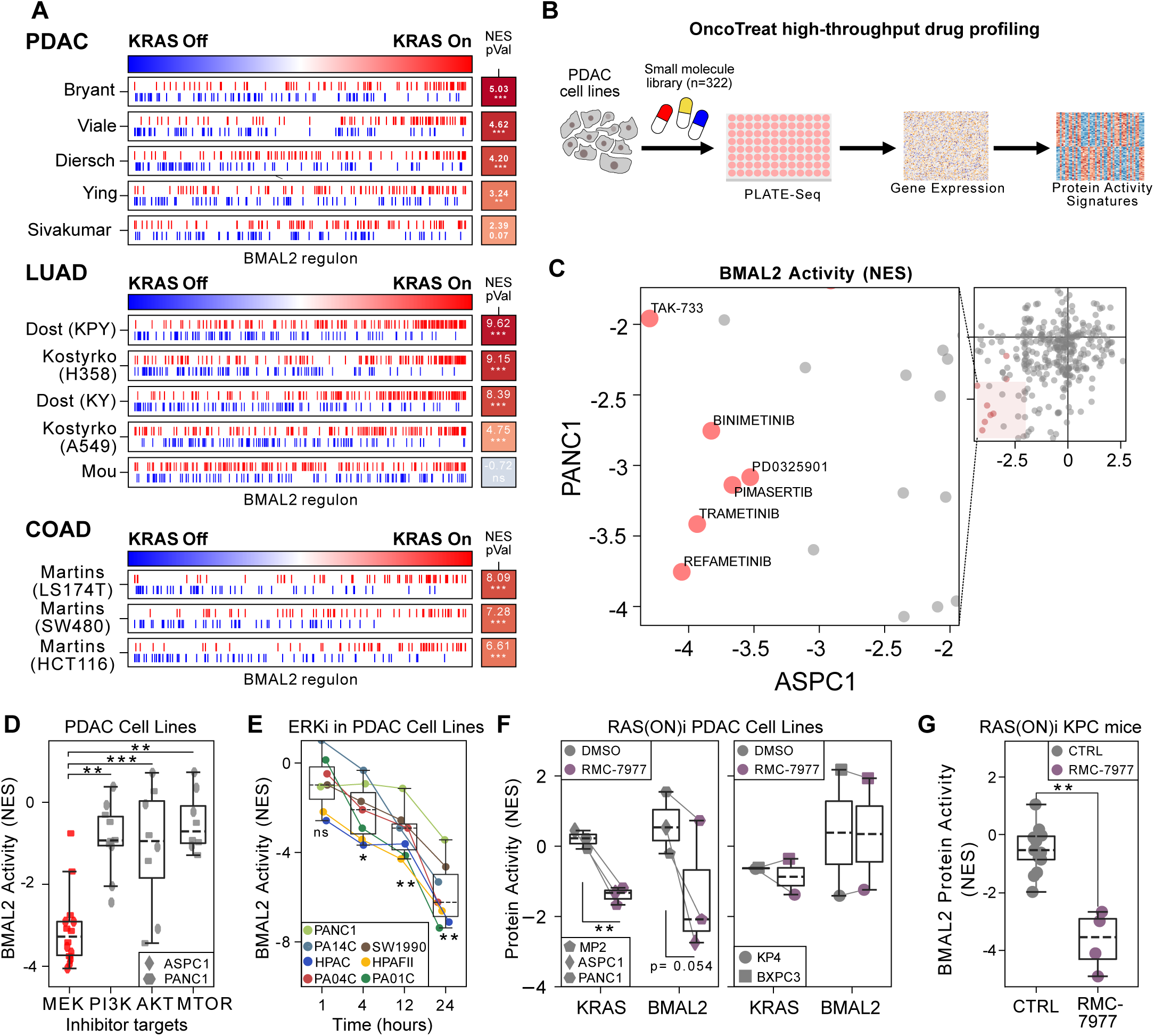
Oncogenic *KRAS* activates BMAL2 via the ERK mitogen-activated protein kinase cascade **(A)** BMAL2 regulon enrichment in the indicated genome-wide response gene expression signatures to the experimental modification of oncogenic *Kras* mutation (Refs. ^44–48^) in PDAC (top panel), lung adenocarcinoma (LUAD, Refs. ^84–86^) and colon adenocarcinoma (COAD, Ref. ^87^). Positive and negative targets, respectively, are represented by red and blue vertical bars, respectively. Normalized enrichment score (NES) and p-values are calculated by two-tailed analytic rank-based enrichment analysis (aREA, p-values are Bonferroni corrected) (**B**) Schematic of the experimental design to study the effect of a library of antineoplastic compounds on regulatory protein activity as part of the OncoTreat framework (Refs. ^49,50^) (**C**) Effects of 322 antineoplastic compounds on BMAL2 activity (NES) in ASPC1 (x-axis) and PANC1 (y-axis) cells. Zoomed area shows compounds with consistent and potent reversal of BMAL2 activity. Red circles mark MEK inhibitors, all other compounds are grey. (**D**) Reversal of BMAL2 activity (y-axis, normalized enrichment score, NES) for the indicated compound classes (x-axis). P-values are derived from pairwise t-tests with post hoc Bonferroni correction. (**E**) Reversal of BMAL2 activity (y-axis, NES) at the indicated time points of treatment with an ERK inhibitor (SCH772984) in 7 PDAC cell lines. (**F**) Inferred changes in KRAS and BMAL2 protein activity in pancreatic cancer cell lines upon RMC-7977 (100nM) treatment for 24h. Statistical significance was determined by a paired, two-tailed t-test, and p-values are indicated where significant (*p < 0.05, **p < 0.01, ***p < 0.001). (**G**) Box plots showing Normalized Enrichment Score (NES) of BMAL2 activity in tumor cells from single-cell RNA sequencing from control and 1-week RMC-7977 treated mice^51^

To evaluate potential regulators of BMAL2 in an unbiased manner, we leveraged results from a recent analysis that presented high throughput RNA-Seq data from two human PDAC cell lines treated with 322 different drugs ^49,50^ (**Figure 2B**). This exercise revealed a strong overrepresentation of MEK inhibitors among the agents most capable of reducing BMAL2 activity in PDAC lines (**Figure 2C**). By contrast, we did not observe effects of this magnitude for inhibitors of other KRAS effector proteins, including PI3K, AKT, and MTOR (**Figure 2D**). Although this screen lacked ERK inhibitors, a reanalysis of experimental expression data from PDAC cells treated with the ERK1/2 inhibitor SCH772984 found decreasing BMAL2 activity over the course of 24 hours after treatment ^47^ (**Figure 2E**).

Finally, to directly test whether RAS inhibition controls BMAL2, we performed RNA-Seq on four human PDAC cell lines treated with RMC-7977, a RAS(ON) multi-selective inhibitor that potently inhibits mutant and wild-type variants of KRAS, HRAS, and NRAS^51,52^. In vitro, we found that BMAL2 activity was significantly decreased upon RAS inhibition in three PDAC lines that were sensitive to RMC-7977 (**Figure 2F**), despite having no impact to BMAL2 expression levels, consistent with a post-translational mechanism of regulation. Interestingly, in two lines with low sensitivity to RAS inhibition due to a BRAF mutation (BXPC3) and MYC amplification (KP4), RMC-7977 treatment did not alter BMAL2 activity. Next, we examined single cell RNA sequencing (scRNA-Seq) data from a collection of pancreatic tumors arising in the *Kras^LSL.G12D/+^*; *Trp53^LSL.R172H/+^*; *Pdx1-Cre^tg/+^* (KPC) mouse model (manuscript in preparation) and applied regulatory network analysis to malignant epithelial cells from mice treated either for one week with RMC-7977 versus controls (**Supplementary Figure S3A**). We found in this *in vivo* experiment that pan-RAS inhibition significantly reduced the activity of BMAL2 in the malignant epithelial cells of PDAC (**Figure 2G**). Together, these data demonstrate that BMAL2 activity is regulated by oncogenic *KRAS* via the RAF/MEK/ERK effector pathway in pancreatic cancer.

### BMAL2 controls hypoxia response targets

BMAL2 belongs to the basic helix–loop–helix PER-ARNT-SIM (bHLH-PAS) family of transcription factors that heterodimerize to drive varied functions including circadian rhythm programs, innate and adaptive immune responses, oxygen-sensing mechanisms, and response to deleterious environmental exposures ^53^. BMAL2 is classically associated with circadian processes, serving as a binding partner for CLOCK ^54,55^, but sequence conservation analysis shows that BMAL1 and BMAL2 are most closely related to ARNT (HIF1B) and ARNT2 (HIF2B), the binding partners of the hypoxia-responsive HIF1A and HIF2A proteins (**Figure 3A**). As hypoxia plays an important role in cancer, we examined the association of activities for each bHLH-PAS family member in our 197 PDAC epithelial profiles with eight publicly available hypoxia transcriptional signatures^56^. We found BMAL2 exhibited the highest average positive correlation (Spearman’s rho 0.46, **Figure 3B**) with hypoxia signatures of any bHLH-PAS family member. Interestingly, neither HIF1A nor HIF1B (ARNT) were strongly correlated with hypoxia signatures in PDAC epithelial samples, despite being among the top correlated PAS family members in laser capture microdissected PDAC stromal samples from the same tumors (N=124, **Figure 3C**). To assess whether BMAL2 activity is associated with hypoxia signatures more broadly in cancer, we built bespoke regulatory networks for multiple tumor types from TCGA datasets and found that BMAL2 was frequently among the most hypoxia-associated bHLH-PAS family members, along with HIF1A and HIF1B (**Figure 3D**). These data suggest that BMAL2 may play an underappreciated role in hypoxia responses across human tumors and a particularly prominent role in the hypoxia response of PDAC.

**Figure 3.**
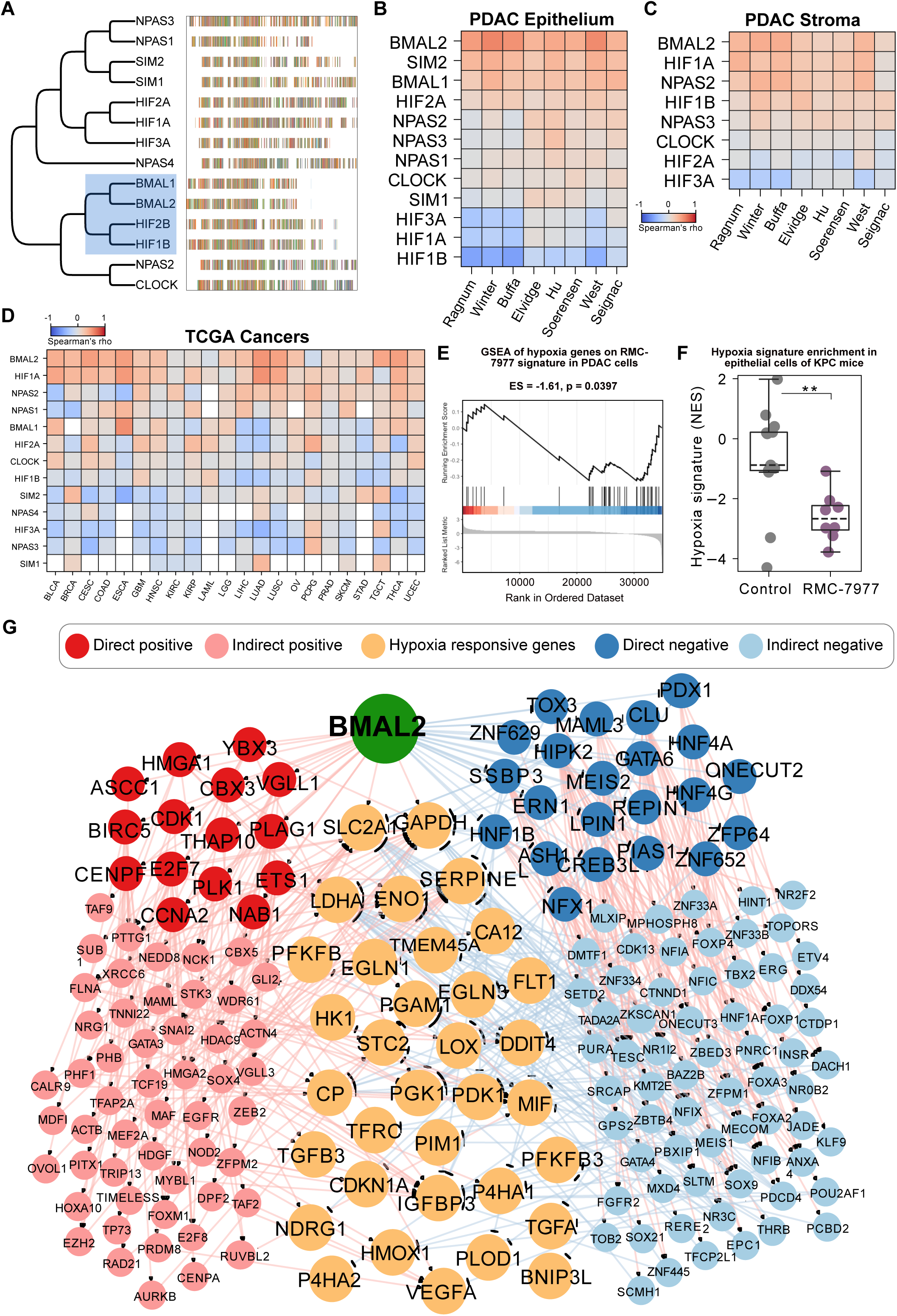
The bHLH-PAS family member BMAL2 controls hypoxia response targets **(A)** Phylogenetic tree illustrating the pairwise distance based on sequence alignment of bHLH-PAS family transcription factors. Blue rectangle highlights the closest family members for BMAL2. Vertical bars represent specific amino acids at the indicated position (1-1300). (**B**) Heatmap of Spearman’s rank correlation between transcriptional hypoxia scores using signatures from the indicated references (x-axis) and epithelial protein activity for bHLH-PAS family members (y-axis). (**C**) Heatmap of Spearman’s rank correlation between transcriptional hypoxia scores using signatures from the indicated references (x-axis) and stromal protein activity for bHLH-PAS family members (y-axis). (**D**) Heatmap of the average Spearman’s rank correlation between eight transcriptional hypoxia scores in the indicated TCGA tumor cohorts (x-axis) and protein activity for bHLH-PAS family members (y-axis). (**E**) Gene Set Enrichment Analysis (GSEA) of hypoxia-related gene signatures upon RAS inhibition in five human PDAC cell lines treated with 100nM of the RAS(ON) inhibitor RMC-7977 for 24 hours. **(F)** Box plots showing Normalized Enrichment Score (NES) of a hypoxia-related gene set^57^ in tumor cells from single-cell RNA sequencing from control and 1-week RMC-7977 treated mice^51^. The normalized enrichment score (NES) and p-value are indicated. (**G**) HIF target genes (Ref. ^57^, orange) controlled directly or indirectly by BMAL2 (green). Indirect control involves both BMAL2’s negative influence on first (dark blue) or second tier (light blue) RP repressing HIF target genes, and positive influence on first (dark red) and second (light red) tier RP activating HIF target genes.

Next, we returned to the RNA-Seq data from RMC-7977 treated PDAC cells and found that RAS inhibition led to a decrease in hypoxia signature scores (**Figure 3E**), concordant with the observed downregulation of BMAL2 activity (**Figure 2F**). This was further validated *in vivo* in the KPC tumor scRNA-Seq dataset, which showed a decrease in hypoxia signature enrichment in malignant epithelial cells following RMC-7977 treatment (**Figure 3F**, Supplementary Figure 3A). To investigate whether BMAL2 could plausibly play a direct role in transcriptionally regulating hypoxia programs, we analyzed the inferred target genes of BMAL2 in the CUMC-E regulatory network and evaluated enrichment for genes canonically controlled by HIF1A ^57^. We found that BMAL2 impacts 33 out of 44 hypoxia genes from a published hypoxia signature ^57^ were regulated in a net positive manner, including metabolic proteins such as SLC2A1 (GLUT1), GAPDH, and LDHA (**Figure 3G**); none of the canonical HIF1A target genes were regulated by BMAL2 in a net negative manner. Together, these analyses support the hypothesis that BMAL2 contributes to the transcriptional regulation of hypoxia genes in PDAC.

### Autochthonous pancreatic tumors are severely hypoxic

The activation of oncogenic KRAS in PDAC provokes a cascade of paracrine signals that suppresses angiogenesis ^58^, resulting in low tumor vascularity and limited perfusion^59^. By inference, these tumors are widely expected to be severely hypoxic, but there are few direct measurements of partial oxygen pressure (pO_2_) in PDAC tissues. Oxygen microelectrode measurements on a small set of human PDAC patients previously indicated the presence of extreme hypoxia (ranging from 0 – 5.3 mmHg) ^60^, but technical concerns limited interpretation ^61^. We therefore measured the oxygenation of autochthonous pancreatic tumors arising in KPC mice – a model system widely utilized for its physiological accuracy to human PDAC. We first measured intratumoral pO_2_ using ultrasound-guided placement of an OxyLite sensor (a gold-standard physical sensor of oxygen) and found that pO_2_ levels were <1mmHg in KPC mouse pancreatic tumors (**Figure 4A, B**), reflecting a setting of extreme hypoxia. This finding was further supported via photoacoustic imaging on KPC pancreatic tumors, which revealed an average hemoglobin saturation of just 17%, significantly lower than in adjacent pancreas tissue (**Figure 4C and Supplementary** Figures 3B-D). Finally, we measured activation of the hypoxia probe pimonidazole ^62^ following its administration to KPC mice respiring normoxic or hypoxic (10% O_2_) air. In these mice, normal tissues showed marker staining only upon exposure to hypoxia whereas PDAC tissues showed equally high levels of staining under both normoxic and hypoxic conditions (**Figure 4D, Supplementary Figure 3B**). Together these experiments demonstrate that autochthonous pancreatic tumors in a physiologically relevant model system naturally exist in a state of extreme hypoxia, underscoring the importance of hypoxia response programs to PDAC biology.

**Figure 4.**
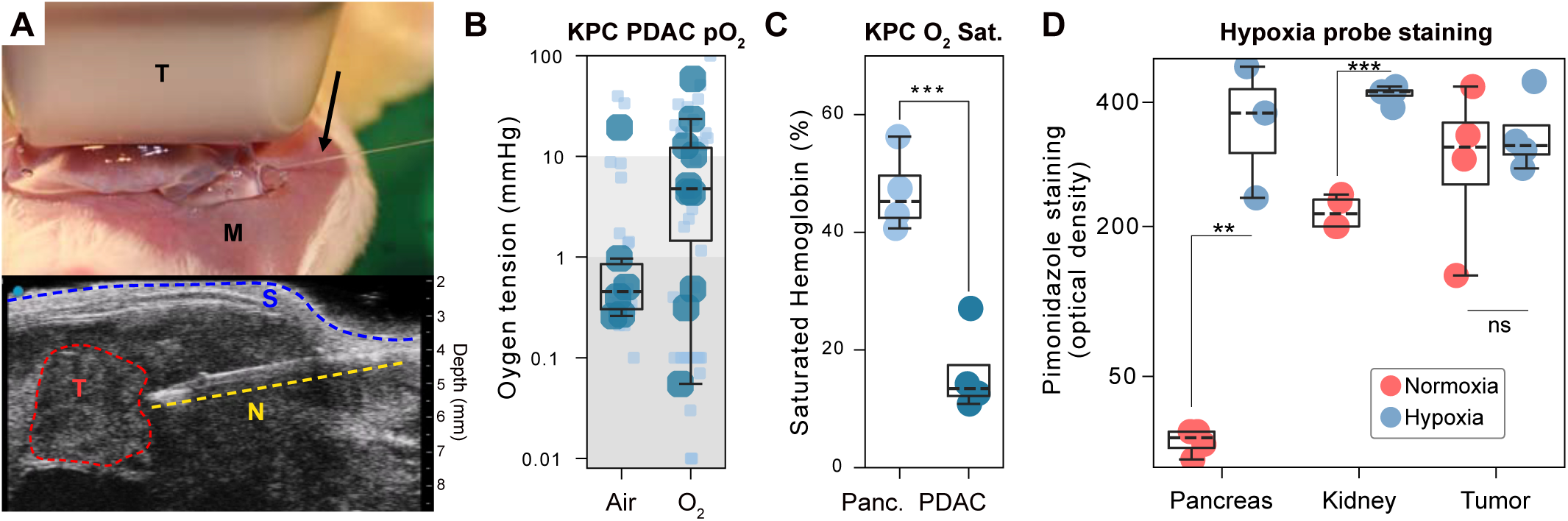
Severe hypoxia in pancreatic tumors is highlighted by multiple methods **(A)** Image (top) shows a KPC mouse (M) being imaged with an ultrasound transducer (T), with percutaneous insertion of the OxyLite probe (arrow). An ultrasound image (bottom) shows the probe (red hashline, offset) extending through the abdominal wall and through the depth of the tumor (outlined in blue). (**B**) Oxygen tension (y-axis) as determined by an OxyLite probe in tumors from KPC mice breathing ambient air or pure oxygen (x-axis). Dark blue circles represent averages per tumor and boxplots illustrate their distribution. Light blue circles represent repeat measurements per tumor at different sites. (**C**) Fraction of saturated hemoglobin (y-axis) in the indicated tissue (x-axis). (**D**) Hypoxia marker pimonidazole staining intensity in the indicated tissues and oxygen conditions. **: p ≤ 0.01, ***: p ≤ 0.001, ns: not significant In boxplots, the box ranges from Q1 (the first quartile) to Q3 (the third quartile) of the distribution and the range represents the IQR (interquartile range). The median is indicated by a dashed line across the box. The “whiskers” on box plots extend from Q1 and Q3 to 1.5 times the IQR.

### BMAL2 drives PDAC cell proliferation and hypoxic metabolism

We next examined the consequences of BMAL2 loss in four human PDAC cell lines (KP4, PANC1, MIAPACA2, and PATU8902) under both normoxic and hypoxic conditions (1% O_2_), using CRISPR/Cas9 genome editing. Under normoxia, loss of *BMAL2* significantly reduced both cell viability, trans-well cell migration, clonogenic growth at low density, and in a scratch invasion assay (**Figure 5A, B** and **Supplementary** Figure 4A-C). We noted that BMAL2 drove the activity of a number of other regulatory proteins, including several several direct positive targets associated with proliferation, including: Cyclin A2 (CCNA2), MET, NRAS, ETS1, BUB1, and AXL (**Figure 5C**). We selected two of these for (AXL and CCNA2) for validation by western blotting in in PDAC cells and confirmed their downregulation in response to *BMAL2* loss (**Figure 5D**). The effects of BMAL2 loss on proliferation were also apparent upon exposure to hypoxia; *BMAL2* knockout reduced cell viability, clonogenic growth, and scratch invasion to a similar degree regardless of oxygen levels (**Figure 5A** and Supplementary Figure 4A-C). However, *BMAL2* loss had a particularly potent effect on trans-well cell migration, synergistically reducing the ability of PDAC cells move through a porous membrane under hypoxia (**Figure 5B**, p_interaction_= 0.004), highlighting a specific contribution of BMAL2 to the hypoxia response of PDAC cells. This is consistent with a deep body of literature linking hypoxia to cell migration and places BMAL2 as a key transcriptional mediator of hypoxia-induced migration ^63,64^.

**Figure 5.**
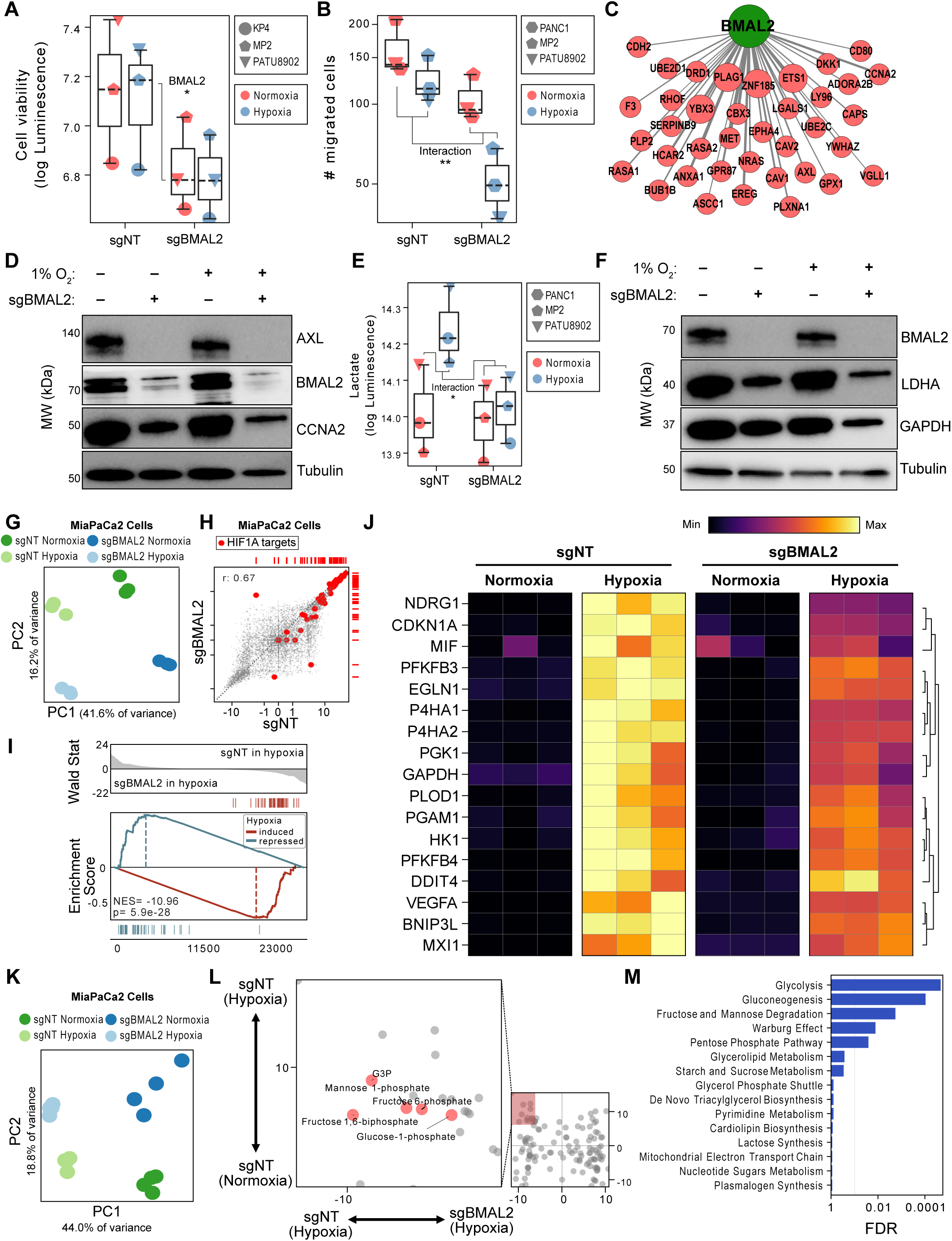
*BMAL2* knockout phenotypes in pancreatic cancer cells are pronounced by hypoxic environments and blunts its transcriptional and metabolic response. **(A)** Cell numbers represented by luminescence in pancreatic cancer cells expressing non-targeting (sgNT) or *BMAL2*-directed (sgBMAL2) sgRNA in the indicated oxygen environment and cell line. (**B**) Number of migrated cells for pancreatic cancer cells expressing sgNT or sgBMAL2 in the indicated oxygen environment and cell line. P-value stems from testing the interaction coefficient between *BMAL2* knockout and hypoxic conditions from a log linear regression model. (**C**) Depiction of a subset of target genes inferred for BMAL2 from the PDAC regulatory network, focused on those associated with proliferation. (**D**) Western Blot for direct targets AXL and CCNA2 in KP4 cells carrying the indicated sgRNA in the indicated oxygen environment (24h). Tubulin was the loading control. (**E**) Extracellular lactate levels after 5 days in pancreatic cancer cells expressing sgNT or sgBMAL2 in the indicated oxygen environment and cell line. P-value stems from testing the interaction coefficient between *BMAL2* knockout and hypoxic conditions from a log linear regression model. *: p ≤ 0.05, **: p ≤ 0.01. (**F**) Western Blot for hypoxia targets LDHA and GAPDH in Patu8902 cells carrying the indicated sgRNA in the indicated oxygen environment (24h). Tubulin was the loading control. (**G**) Principal component analysis (PCA) based on gene expression of the indicated cell line carrying sgNT or sgBMAL2 in hypoxic or normoxic environments, respectively. (**H**) Scatter plot illustrating the relationship of a genome-wide transcriptional hypoxia signature found in the indicated cell lines carrying sgNT (x-axis) and sgBMAL2 (y-axis) sgRNA, respectively. Red circles mark a set of HIF1A reporter genes described previously (Ref. ^57^) (**I**) 2-tailed GSEA of the top 100 transcripts induced (red) and repressed (blue) by hypoxia in PDAC cells on a gene expression signature between hypoxic sgBMAL2 cells (left) and sgNT cells (right). (**J**) Heatmap illustrating HIF1A reporter gene expression (Ref. ^57^) for a subset with a differential hypoxia response between cells carrying sgNT or sgBMAL2, respectively. (**K**) PCA based on metabolite abundances in sgBMAL2 vs sgNT MP2 cells in hypoxic or normoxic environments, respectively. (**L**) Differential abundance signatures of 230 metabolites in MP2 cells comparing the effects of hypoxia treatment (y-axis) and knockout of BMAL2 (x-axis). We focused on metabolites that were upregulated under hypoxia more in BMAL2 wild-type cells than in BMAL2 knockout cells (inset). (**M**) shows metabolite sets that are overrepresented among this group.

A key component of hypoxic metabolism in all cells is the production and secretion of the glycolytic product lactate ^65^. We examined the impact of *BMAL2* knockout on lactate secretion in PDAC cells after five days in culture. As expected, exposure to hypoxia significantly increased extracellular lactate levels of control cells expressing a non-targeting sgRNA (sgNT). However, upon loss of *BMAL2*, hypoxia no longer increased lactate secretion (**Figure 5E**, p_interaction_ = 0.025), indicating a strong reliance of PDAC cells on BMAL2 function to facilitate the hypoxia-induced shift to glycolytic metabolism. Indeed, Western blots for LDHA and GAPDH, two glycolysis proteins that were identified in the PDAC regulatory network as indirect targets of BMAL2, found that both were reduced in response to *BMAL2* knockout. Moreover, expression of Lactate Dehydrogenase A (LDHA), which is directly responsible for cellular lactate production in the final step of glycolysis, was decreased in *BMAL2* null PDAC cells after 24 hours under hypoxia (**Figure 5F**), further supporting the role of BMAL2 in driving hypoxic metabolism.

To better understand the molecular consequences of BMAL2 loss in PDAC cells, we performed transcriptomic profiling of KP4 and MIAPACA2 cells from hypoxic and normoxic environments. Unsupervised clustering of gene expression profiles showed that hypoxia exposure and *BMAL2* knockout dominated the global variance in both cell lines (**Figure 5G** and **Supplementary** Figure 4D). In both *BMAL2* wild type and knockout lines, exposure to hypoxia induced transcriptional programs enriched for HIF1A target genes (**Figure 5H** and **Supplementary** Figure 4E), indicating that BMAL2 knockout cells are still capable of mounting a transcriptional response to hypoxia. However, the magnitude of their response to hypoxia was significantly blunted upon loss of BMAL2 in both cell lines, as demonstrated by directly comparing the enrichment of hypoxia signatures in sgNT versus sgBMAL2 cells (**Figure 5I**). Moreover, we found that 19 out of 44 hypoxia signature genes ^57^ were significantly less activated by hypoxia upon BMAL2 deletion (FDR_interaction_ < 0.05, **Figure 5J**).

Next, we examined whether the transcriptional programs altered by BMAL2 loss impacted the metabolic programs induced by hypoxia in PDAC cells. We used an LC/MS metabolomic panel to quantify ∼230 metabolites from *BMAL2* wild type or knockout PDAC cells exposed to normoxia or hypoxia (**Figure 5K**). Guided by our transcriptional findings, we focused on metabolites that increased in hypoxic conditions but to a lesser degree in *BMAL2* knockout cells compared to wildtype cells (**Figure 5L**). Among these, metabolite set analysis found significant overrepresentation of metabolites associated with glycolysis and related metabolic pathways (**Figure 5M**) including fructose 6-phosphate, fructose 1,6-bisphosphate, and glyceraldehyde 3-phosphate (G3P). Together these results that BMAL2 broadly sculpts the transcriptional responses of PDAC cells to modulate the metabolic responses to hypoxia.

Finally, to determine whether the phenotypes we observed in cultured PDAC cells ultimately impact tumor growth, we assessed the effect of BMAL2 knockout *in vivo* using two human PDAC cell line-derived xenografts (CDX) implanted orthotopically in the pancreas of immune-deficient mice (**Figure 6A**). First, in orthotopic PANC1-derived tumors, we found that loss of BMAL2 significantly reduced tumor growth rates, as measured by longitudinal 3D high resolution ultrasound (**Figure 6B,C**). We then repeated the experiment using KP4 cells and found that BMAL2 had a profound impact on tumor engraftment, with 0 of 12 implanted mice exhibiting tumors after 8 weeks, compared to 8 of 12 (66.67%) from sgNT expressing cells (**Figure 6D**). We conclude that BMAL2 is a master transcriptional regulator of PDAC malignancy that both promotes cell proliferation and drives the transcriptional and metabolic responses to hypoxia.

**Figure 6.**
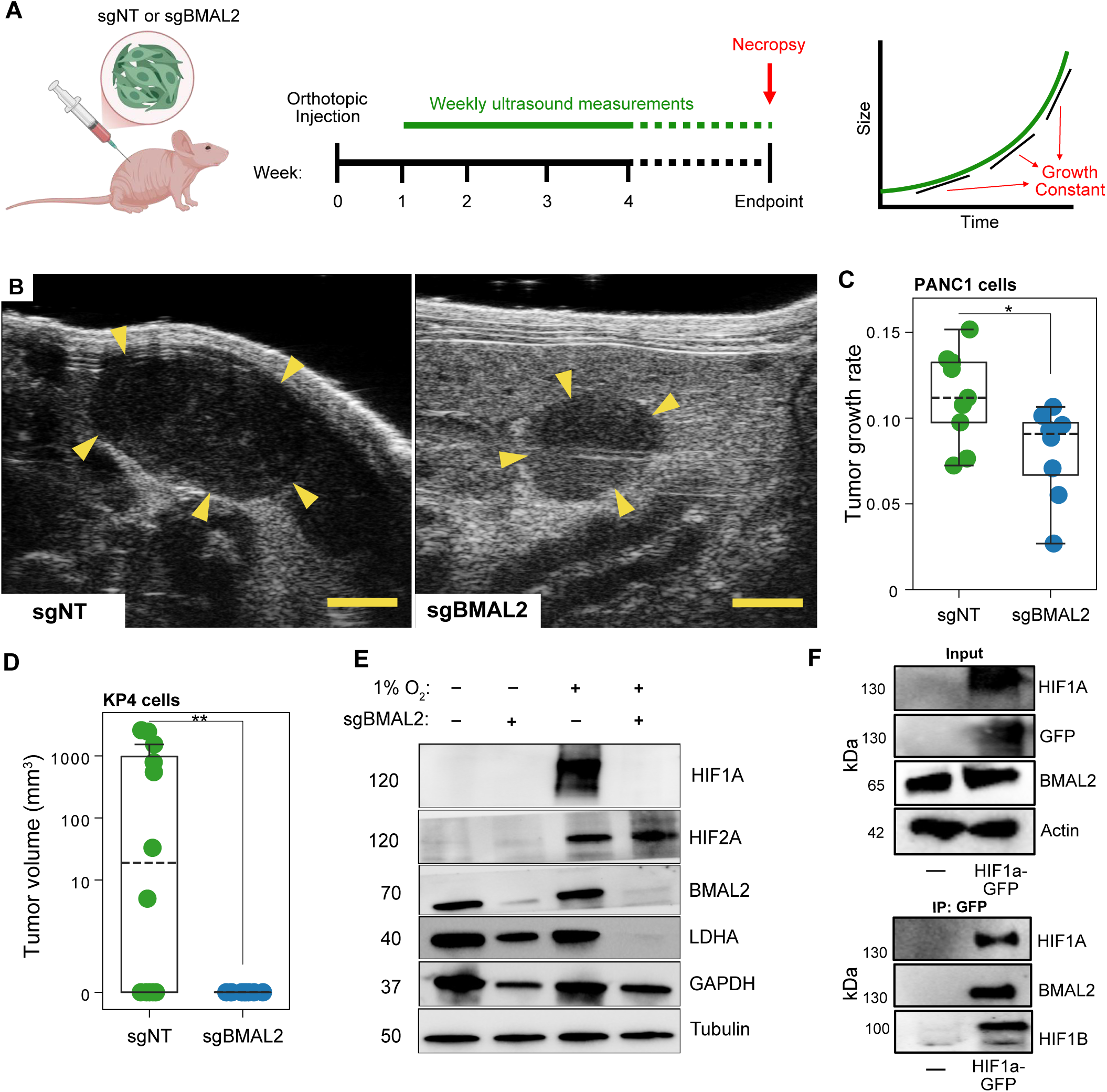
HIF1A interacts with BMAL2 and depends on it for stabilization in pancreatic cancer cells **(A)** Experimental design of the *in vivo* experimental setup. Immunodeficient NOD/SCID mice were orthotopically injected with PANC1-Cas9 or KP4-Cas9 cells carrying non-targeting control (sgNT) or sgBMAL2. Tumor growth was monitored weekly using longitudinal, high-resolution 3D ultrasound imaging. At endpoint, tissue was collected and processed for analysis. (**B**) Representative ultrasound images of orthotopic tumors derived from PANC1 sgNT and sgBMAL2 cells. Yellow arrowheads indicate tumor boundaries. Scale = 2mm. (**C**) Box plot showing tumor growth rates (y-axis) in tumors carrying the indicated sgRNA (x-axis) in PANC1 cells. Groups were compared by two-tailed Mann-Whitney U test. (**D**) Box plot showing tumor volumes after 8 weeks in sgNT and sgBMAL2 KP4 cells. Groups compared by two-tailed Mann-Whitney U test. (**E**) Western Blot for HIF1A, HIF2A, BMAL2, LDHA and GAPDH in PANC1 cells carrying the indicated sgRNA in the indicated oxygen environment (24h). Tubulin was the loading control. (**F**) Input Control: Total cell lysates were subjected to Western blotting to verify the expression levels of HIF1A, GFP and BMAL2. Co-IP. For Co-immunoprecipitation (Co-IP), GFP-tagged HIF1A was immunoprecipitated from cell lysates (*top panel*), and the presence of interacting proteins was assessed by Western blotting using antibodies against HIF1A, BMAL2, and HIF1B (ARNT) (bottom *panel*).

### BMAL2 reciprocally regulates the stability of HIFα proteins

The classical cellular responses to hypoxia are mediated by the hypoxia inducible factors HIF1α and HIF2α ^66^, which are stabilized in response to low oxygen levels. The HIFα proteins are first stabilized through loss of (oxygen-dependent) proteasomal degradation, followed by further stabilization through heterodimerization with the transcription factor HIF1β ^67^. To assess the impact of BMAL2 on HIF-dependent hypoxia regulation, we performed western blots for HIF1α and HIF2α on BMAL2 knockout and wild type PDAC cells, cultured in normoxia and hypoxia (**Figure 6E**). Strikingly, knockout of BMAL2 fully prevented the stabilization HIF1α under hypoxia in four different PDAC cell lines, consistent with the observed decrease in LDHA expression. In contrast, HIF2α accumulated to even higher levels in BMAL2 knockout cells than in wild-type cells, suggesting that BMAL2 can serve as a switch between HIF1α- and HIF2α-dependent modes of hypoxia response.

Given the evolutionary conservation of BMAL2 and HIF1β, we considered whether BMAL2 contribute to HIF1α stabilization by serving as a heterodimerization partner. To test this, we transfected HEK293 cells with a GFP-tagged HIF1α construct and performed coimmunoprecipitation and western blotting (**Figure 6F**). In addition to detecting the canonical partner HIF1 we were able to detect both endogenous BMAL2 and HIF1β in complex with HIF1α, suggesting that BMAL2 may play a direct role in regulating the stability of HIFα family members. In summary, we find that BMAL2 serves as a RAS-dependent regulator of hypoxia transcriptional programs that drive PDAC malignancy.

## Discussion

The stability of key genetic mutations throughout PDAC progression, from precursor lesions to metastasis, suggests that while these mutations initiate the disease, additional, non-genetic mechanisms must drive the dynamic changes in tumor behavior and aggressiveness observed in later stages. Here we utilized regulatory network analysis to explore the landscape of non-genetic regulators of PDAC. Anchoring our expression profiles to histopathological annotation, we find that epithelial differentiation state is closely mirrored by transcriptional regulatory programs. The availability of profiles from benign precursors contextualizes our observations of malignant samples. For example, oncogenic properties have frequently been ascribed to transcription factors, such as GATA6, that are “overexpressed” in the (well-differentiated) Classical subtype, relative to other PDAC tumors ^17,26^. However, GATA6 activity was significantly higher in benign precursors than in low-grade tumors. We infer that the comparatively high activity of GATA6 in low-grade tumors reflects an incomplete loss of differentiation state—consistent with functional characterizations ^68,69^—that point to a largely tumor-suppressive role for GATA6. We find a similar pattern of downregulation in PDAC for the majority of pancreatic transcription factors, including PDX1, HNF1B, and SOX9. These data prompt a reevaluation of their strict classification as ‘drivers’ of low-grade, classical tumors, especially considering that loss of differentiation in high-grade tumors is associated with worse prognosis ^70,71^. We urge caution in attempts to target GI transcription factors therapeutically, at any stage of disease, and would instead favor strategies that target consistent RPs of multiple malignant phenotypes.

While virtually all aspects of PDAC biology are influenced by activating mutations in *KRAS*, their association to histopathological phenotypes is limited ^72,73^. We find KRAS activity is lowest in low-grade precursor lesions, despite the high prevalence of activating *KRAS* mutations ^1^ in PanIN. With the progression to PDAC and eventual loss of differentiation ^74^, KRAS activity increases steadily. Among the 2,211 regulatory proteins we measured, this pattern was most strongly correlated with the activity of BMAL2, a transcription factor that is largely undescribed in pancreatic cancer. Our pharmacologic perturbation data, particularly the treatment of PDAC cell lines with RAS, MEK and ERK inhibitors, demonstrates that BMAL2 activity is effectively regulated by RAS/MAPK signaling. This finding places BMAL2 in company with well-validated downstream effectors of the RAS-MEK-ERK cascade such as MYC ^44,75^ and ETS1 ^76^ (the latter of which is a direct target of BMAL2 in our regulatory network).

Although BMAL2 is classically associated with circadian rhythm function ^53^, our results show that BMAL2 is a critical regulator of hypoxia responses. As we demonstrate, the hypovascularity of PDAC ^59^ results in a state of profound hypoxia, begging the question of how these tumors can survive and thrive in such an adverse environment. Certainly, severe hypoxia can confer several advantageous phenotypes, including immunosuppression, inflammation, invasiveness, and EMT, and an associate between RAS signaling and glycolysis has long been apparent from cell culture studies ^44,77,78^. However, the precise mechanisms by which PDAC cells survive such an extreme environment have remained cryptic given that tumor-cell specific deletion of Hif1a in KPC mice counterintuitively accelerates PDAC progression ^79^. Our current findings on BMAL2 demonstrate that KRAS mutation is directly linked both to the paracrine suppression of angiogenesis in PDAC^58^ and to a cell-autonomous regulatory program that enables survival in the resulting hypoxic microenvironment.

It has long been appreciated that HIF1α and HIF2α are differentially regulated through an unknown mechanism and that they drive distinct metabolic and transcriptional responses ^80,81^. The possibility that BMAL2 can serve as a functional switch between HIF1α- and HIF2α-dependent hypoxic responses provides a potential answer to the long-standing question of how these proteins are differentially regulated. With multiple HIF-targeted drugs entering clinical development ^82^, it will be critical to understand in which settings these proteins serve as critical dependencies and for which they are dispensable. Moreover, given that BMAL2 knockout mice are viable into adulthood with only modest physical phenotypes ^83^, we anticipate that parallel chemical approaches for targeting BMAL2 and other bHLH-PAS proteins may enable yield novel therapeutic strategies for targeting the hypoxia response of a broad range of cancers.

## Data Availability Statement

The data generated in this study are available upon request from the corresponding author.

## Ethics Statement

All mouse studies performed were approved by the Columbia University Irving Medical Center (CUIMC) Institutional Animal Care and Use Committee (IACUC). All work using patient samples was performed with approval from the CUIMC Institutional Review Board (IRB).

## Acknowledgments

K.P.O was supported by the NCI (1U01CA274312), by a Career Development Award from the Pancreatic Cancer Action Network, and a Clinical Translational Research Award from the Lustgarten Foundation for Pancreatic Cancer Research. C.A.L was supported by the NCI (R01CA244931). A.C.G received support from the Charles H. Revson Senior Postdoctoral Fellowship in Biomedical Science (Grant No. 22-22). K.J. received research supported by New Hampshire-INBRE through an Institutional Development Award (IDeA), P20GM103506, from the National Institute of General Medical Sciences of the NIH. H.C.M received support from a Mildred Scheel Postdoctoral Fellowship (Deutsche Krebshilfe) and the KKF Clinician-Scientist program (TUM). This work utilized resources from the CU-DLDRC (P30 DK132710) and the Columbia University CCSG (5P30CA013696), including the Genomics, OPTIC, Database, and Molecular Pathology Shared Resources. RMC-7977 material was provided by Revolution Medicines under the terms of a sponsored research agreement.

## Author contributions

H.C.M. and K.P.O conceived the project. H.C.M. started the project and performed the LCM dataset and the computational analyses. A.C.G. designed, performed and oversaw the experimental analyses and contributed to the computational analyses. S.H., C.C., C.F.P, S.A.S, A.A., L.Z., T.L., I.S., D.R.R., K.S., and J.G. contributed to experimental execution, data acquisition and data generation. G.A.M. and W.W. aided in the curation of human outcomes data. A.I. reviewed histopathology for pancreatic specimen used in the CUMC cohort. H.C.M, A.C.G., M.A.B and K.P.O. wrote the manuscript with feedback and editing from R.M.S., K.J., C.A.L, Y.S and A.C.

## Methods

### Patient population and samples generation

#### Patient population

Freshly frozen tissue samples were obtained from patients who underwent surgical resection at the Pancreas Center at Columbia University Medical Center as described previously ^11^. The clinical data of these patients are shown in Supplementary Tables S1 and S2. Before surgery, all patients had given surgical informed consent, which was approved by the institutional review board. Immediately after surgical removal, the specimens were cryopreserved, sectioned, and microscopically evaluated by the Columbia University Tumor Bank (IRB AAAB2667). Suitable samples were transferred into OCT medium (Tissue Tek) and snap-frozen in a 2-methylbutane dry ice slurry. The tissue blocks were stored at −80°C until further processing. H&E stained sections of frozen PDAC samples from the Tumor Bank were initially screened to confirm the diagnosis and overall sample RNA quality was assessed by the Pancreas Center supported Next Generation Tumor Banking program using gel electrophoresis, with samples exhibiting high RNA quality utilized for subsequent analyses.

#### Laser Capture Microdissection (LCM), RNA sequencing, and gene expression quantification

LCM-RNA-Seq was performed as described previously ^11,88^. Briefly, Cryosections of OCT-embedded tissue blocks were transferred to PEN membrane glass slides and stained with cresyl violet acetate. Adjacent sections were H&E stained for pathology review. Laser capture microdissection was performed on a PALM MicroBeam microscope (Zeiss), collecting at least 1000 cells per compartment. RNA was extracted and libraries were prepared using the Ovation RNA-Seq System V2 kit (NuGEN). Libraries were sequenced to a depth of 30million, 100bp, single-end reads on an Illumina HiSeq 2000 or 4000, respectively, platform. Reads and transcripts per million (TPM) were estimated for each transcript using the transcript sequences from the GENCODE Release 34 (GRCh38.p5) and the Salmon software (v1.3.0). Counts and TPM were summarized at the gene level by summing up the transcript values for each corresponding gene.

### Computational methods

#### Assembly of a PDAC regulatory model and network analysis

A cell regulatory network for pancreatic carcinogenesis (CUMC-E interactome) was reverse-engineered by ARACNe-AP ^7^ using 242 epithelial LCM-RNA-Seq gene expression profiles. Genes with detection rates at 10 counts below 25% in all of the examined conditions (PanIN, IPMN, and PDAC) were removed and the variance was stabilized by fitting the dispersion to a negative binomial distribution as implemented in the *DESeq2* R package ^89^. ARACNe was run with standard settings (using data processing inequality (DPI), with 100 bootstrap iterations using human gene symbols mapping to a set of 1665 transcription factors and 1025 transcription cofactors as described by AnimalTFDB 3.0 ^90^. For the signaling network, a set of 3370 signaling-pathway-related genes was considered which were annotated in the GO Biological Process database as GO:0007165—‘signal transduction’ and in the GO Cellular Component database as GO:0005622—‘intracellular’ or GO:0005886—‘plasma membrane’. Thresholds for the tolerated DPI and mutual information P value were set to 0 and 10–8, respectively. Using the strategy outlined above, we also generated a stromal regulatory network leveraging a set of 159 stromal LCM gene expression profiles (124 PDAC, 19 PanIN, 12 IPMN). Cytoscape v3.7.1 ^91^ was used to illustrate subnetworks of relevant regulatory proteins. The CUMC-E interactome is available as R object from https://doi.org/10.6084/m9.figshare.13160078.v2.

#### Master Regulator Analysis and inference of virtual protein activity

The enrichment of each regulatory protein’s regulon in the progression and dedifferentiation signature, respectively, was inferred by the MARINa algorithm as implemented in the *msviper* function from the *viper* R package ^28,29,92^. Statistical significance was estimated by permuting the sample labels uniformly at random 1,000 times. For single-sample analysis including precursor and tumor samples, unsupervised gene expression signatures were computed by a z-score transformation of the variance-stabilized data. This was performed gene-by-gene, by first subtracting the mean expression level across all samples and then dividing by its standard deviation. Relative protein activity was then inferred for each sample with the VIPER algorithm. For a patient-based approach including the assessment of whether significant dysregulation of an individual regulatory protein occurs in a given tumor, single sample gene expression signatures were computed for each primary sample by subtracting the mean of all precursor samples (n = 45) and dividing by their standard deviation. RP activity was then inferred by VIPER analysis of each PDAC gene expression signature. P-values were estimated by the analytical approximation implemented in the aREA algorithm, which is virtually equivalent to estimations obtained by permuting the genes in the signature uniformly at random ^29^. P-values were corrected to account for multiple hypothesis testing by the Benjamini-Hochberg method.

#### Gene set enrichment analysis and scoring

One-tail gene set enrichment analysis was implemented as described ^93^. Two-tailed gene set enrichment analysis and enrichment of individual ARACNe regulons were carried out using analytic rank-based enrichment analysis (aREA) ^29^. Functional annotation of ARACNe-derived regulons was carried out by testing the overrepresentation of HALLMARK gene sets (MSigDb v6.0) among all target genes of a given regulatory protein with the gene universe set to all unique genes in the CUMC-E interactome (n=18658) using a two-tailed Fisher’s Exact test.

#### Differential gene expression

Genome-wide differential gene expression analysis was generally calculated using the *DESeq2* ^89^ R package for RNA-Seq count data and the *limma* R package ^94^ for microarray data. For comparisons including both RNA-Seq and microarray data, differential gene expression for count data was repeated using the voom-limma framework for the sake of consistency with the microarray analysis.

#### Survival signature and assessment of prognostic relevance

In an unbiased approach to study the association of gene expression and protein activity with patient outcome, a genome-wide survival signature was computed by fitting a multivariate Cox Proportional Hazards model (CPHM) accounting for the age at diagnosis and the continuous, normalized expression of a respective gene using the *survival* R package ^95^. Next, we extracted the ensuing Wald statistic of the coefficient for gene expression with higher values corresponding to higher hazard ratios and vice versa and applied MARINa with a gene permutation null model to this survival signature to infer regulatory proteins controlling the survival signature.

#### Effect size meta-analysis

The effect size (i.e. log_2_ fold change) and its standard error for BMAL2 were extracted from the respective genome-wide differential expression analysis of a total of 10 studies where global expression in normal vs. primary tumors and low-grade vs. high-grade tumors, respectively, was contrasted. Similarly, the coefficient (log hazard ratio) and its standard error for the upper tertile of BMAL2 expression and activity, respectively, were extracted from a CPHM in cohorts with available time-to-event data. Meta-analysis was carried out using the *metafor* R package ^96^. Both random and fixed effect models were fit using the *rma* function (method = “REML” and method = “FE”).

#### Molecular subtyping

Moffitt classes were determined for 197 primary PDAC LCM-RNA-Seq profiles as described previously ^32^. Briefly, using the 50 (47 with a unique match in our data) tumor-specific transcripts from Moffitt et al., we applied consensus clustering to our mRNA cohort with Euclidean distance and the partitioning around medoids (PAM) algorithm, seeking and reproducing two clusters of both genes and samples.

#### PAS family sequence alignment

Amino acid sequences for bHLH-PAS family members were retrieved using functionality from the Universal Protein Resource Knowledgebase ^97^. After sequence alignment pair-wise distances determined based on sequence identity. Results were illustrated using the *ggtree* and *msatools* R packages.

#### Hypoxia scoring

Using tumor epithelial expression data, hypoxia scores were calculated by using mRNA-based signatures as described previously ^56^. For each gene in each of eight signatures, TPM were extracted and if a tumor’s abundance value exceeded the median across all tumors, +1 was added to its hypoxia score while −1 was added otherwise.

### External data

#### Human PDAC cohorts

For the TCGA-PAAD cohort, raw count data were retrieved from the GDC Data portal for 149 patients described previously ^38^ and the variance was stabilized as described above. For the ICGC-PACA-AU cohort described previously ^40^, normalized gene expression data for 96 patients were provided by the authors in Suppl. Table 2. Microarray data of primary PDAC specimen from Collisson et al. ^26^ (n = 27), Moffitt et al. ^32^ (n = 252), and Puleo et al. ^39^ (n = 309) were retrieved from GSE17891, GSE71729 and E-MTAB-6134, respectively. Studies containing expression data from both normal pancreas and PDAC were processed and analyzed as described previously ^98^.

#### TCGA Pan-Cancer Atlas

Processed clinical and expression data were retrieved from the Genomic Data Commons Pan-Cancer homepage (https://gdc.cancer.gov/about-data/publications/pancanatlas). Information on sample types was added using TCGA sample barcodes. For regulatory network assembly, only tumor types with at least 100 RNA-Seq samples were retained and networks were generated using ARACNe as described above for CUMC samples.

#### Gene perturbation signatures

Raw microarray data from a mouse model harboring a dox-inducible oncogenic *Kras^G12D^* were retrieved from GSE32277 ^44^ and GSE58307 ^45^ and processed using the *affy* and *gcrma* R packages. Processed microarray data from normal murine ductal cells in which oncogenic *Kras^G12D^*was turned on using adenoviral Cre recombinase were retrieved from GSE89846 and E-MTAB-2592 ^46,48^. Raw RNA-Seq data from human PDAC cell lines stably transfected with shRNA targeting *KRAS* were retrieved from the European Nucleotide Archive under accession PRJEB25797 ^47^ and quantified using the pipeline outlined above for CUMC samples. RNA-Seq counts were analyzed using the voom-limma framework implemented in the *edgeR* and *limma* R packages.

#### ChIP-Atlas

We identified ChIP-Seq experiments concerning transcription factors in a cellular context pertinent to pancreatic ductal adenocarcinoma via the ChIP-Atlas ^23^ website. The results of this search are listed in Supplementary Table S3. For each experiment and transcription factor, we tested whether ARACNe-inferred targets of the respective transcription factor were overrepresented among ChIP-Seq inferred targets (10kb window) using a Fisher’s Exact test with subsequent adjustment of p-values using the Benjamini-Hochberg method.

### Experimental methods

#### Cell culture

KP4, PANC1, MIAPACA2 (MP2), PATU8902, ASPC1 cell lines were obtained from ATCC and tested negative for mycoplasma infection. Cells were maintained under standard conditions at 37°C and 5% CO_2_ using manufactured cell media conditions.

#### Genome editing and transfection protocol

The sgRNA (small-guide RNA) for knocking out *BMAL2* as well as a non-targeting (NT) sequence were purchased from GenScript (Piscataway, NJ) using the pLentiGuide-Puro vector as a backbone. Human PDAC cells were first infected with a pLentiCas9-Blast vector (Addgene, #52962) for a constitutive Cas9 expression and selected with Blasticidin (AG Scientific, #B-1247-SOL). Cells expressing Cas9 were then infected with selected virus carrying sgBMAL2 and sgNT, respectively, using the lentiviral protocol according to the manufacturer’s instructions.

#### Viability, lactate and migration assays

Cells were seeded at 3×10^3^ cells per well in 96 well plates and incubated for five days either under normoxia or moved to the hypoxia chamber with 1% O_2_ (StemCell, #27310). Lactate assay was performed using 5 µl of media and the luminometric Lactate-Glo assay kit (Promega, #J5021) according to the manufacturer’s protocol. To measure cell viability, Alamar Blue reagent (Bio-rad, # BUF012) was added to the culture media for 4 h, and absorbance was determined at 570 and 600 nm using a Varioskan LUX Multimode Microplate Reader (Thermo, #3020-80389). For the migration assay, PDAC cells carrying sgNT or sgBMAL2, respectively were seeded (1×10^5^) into transwell membrane inserts in serum-free culture media (5 µm pore, Corning #3412) and regular media was added to the lower chamber. Cells were incubated for 24h in regular conditions. After the incubation time, plates were incubated either under normoxia (37°C, 5% CO_2_, 20%O_2_) or moved to the hypoxia chamber (37°C, 5% CO_2_, 1%O_2_) for 14 hours and cells that migrated across the membrane were fixed and stained with crystal violet and counted under the microscope. For the dose-response assays, KP4 cell line was tested for sensitivity to RMC-7977 in quintuplicates with serial dilutions of RMC-7977 (top concentration of 10 µM) or DMSO. Cells were incubated for 72 h prior to measurement using Alamar Blue. A total of 3 biological replicates were done. Growth percentage was calculated by normalizing drug-treated values to DMSO control, which was set to 100%. Mean ± s.d. was plotted for each tested dilution.

#### Colony formation assays

KP4 cells carrying non-targeting control (NT) or sgBMAL2 ( 10^3^ cells/well) were cultured in 6-well plates at 37 °C under normoxia or hypoxia conditions. After ten days, cells were stained with crystal violet solution and scanned.

#### Wound healing assay

KP4 cells carrying non-targeting control (NT) or sgBMAL2 were seeded on the 6-well plate. Cells were grown into monolayer and manual scratching with a 200 μl pipette tip. Cells were rinsed with PBS and incubated at 37 °C in serum-free media for 24h under normoxia or hypoxia conditions. Photographs of the wounded areas were taken by phase-contrast microscopy.

#### Immunoblotting

Cell pellets were lysed with RIPA lysis buffer (Cell Signaling, #9806S) and protein concentrations were determined by BCA protein assay (Thermo Scientific, #23227) according to the manufacture’s protocol. Proteins were separated on Mini-PROTEAN TGX gels (Bio-Rad, #4561093) and transferred to nitrocellulose membrane (Bio-Rad, #1704156). Membranes were incubated in blocking buffer (5% BSA, 0.1% Tween-20, 10 mM Tris at pH 7.6, 100 mM NaCl) for 1 hour and then with primary antibody overnight at 4 °C according to the antibody datasheet. Antibodies used: HIF1A (1:1000, Cell Signaling, #36169S), beta-Actin (1:1000, Cell Signaling, #4970S), HIF2A (1:1000, Cell Signaling, #7096S), BMAL2 (1:500, Abcam, #ab221557), LDHA (1:1000, Cell Signaling, #3582S), GAPDH (1:1000, Cell Signaling, #2118S), AXL (1:1000, Cell Signaling, #8661), CCNA2 (1:1000, Cell Signaling, #67955) and Tubulin (1:1000, Cell Signaling, #2146S). Anti-rabbit-HRP (Cell Signaling, #7074S) conjugated antibodies was used to detect the desired protein by chemoluminescence with ECL (Bio-Rad, #170-5061).

#### Metabolomics

Metabolites were extracted from cell pellets by adding 1mL of ice-cold 80% MeOH / 20% H_2_O. Samples were vortexed and incubated on dry ice for 10 minutes, centrifuged at 12,000g, with subsequent extraction of the supernatant. Supernatant was volume normalized across sample groups and then concentrated using a SpeedVac Vacuum Concentrator (model: SPD1030), and reconstituted in 50uL of 50% MeOH / 50% H_2_O. Samples were run on a tandem liquid-chromatography mass spectrometry (LC/MS) set up consisting of an Agilent 1290 Infinity II LC and Agilent 6470 Triple Quadrupole (QqQ) mass spectrometer. The method and column parameters were as follows: Solvent A: 97% water and 3% methanol 15 mM acetic acid and 10 mM tributylamine (pH of 5). Solvent C: 15 mM acetic acid and 10 mM tributylamine in methanol. Solvent D for washing is acetonitrile. LC system seal washing solvent: 90% water and 10% isopropanol. Needle wash solvent: 75% methanol and 25% water. GC-grade Tributylamine 99% (ACROS ORGANICS), LC/MS grade acetic acid Optima (Fisher Chemical), InfinityLab Deactivator additive, ESI–L Low concentration Tuning mix (Agilent Technologies), LC-MS grade solvents of water, and acetonitrile, methanol (Millipore), isopropanol (Fisher Chemical). Column: Agilent ZORBAX RRHD Extend-C18, 2.1 × 150 mm and a 1.8 µm and ZORBAX Extend Fast Guards for ultra high-performance liquid chromatography (UHPLC). LC gradient profile: 0.25 mL/min, 0–2.5 min, 100% A; 7.5 min, 80% A and 20% C; 13 min 55% A and 45% C; 20 min, 1% A and 99% C; 24 min, 1% A and 99% C; 24.05 min, 1% A and 99% D; 27 min, 1% A and 99% D; at 0.8 mL/min, 27.5–31.35 min, 1% A and 99% D; at 0.6 mL/min, 31.50 min, 1% A and 99% D; at 0.4 mL/min, 32.25–39.9 min, 100% A; and at 0.25 mL/min, 40 min, 100% A. Column temperature was kept at 35°C, samples were at 4°C and the injection volume was 2 µL per sample. The 6470 Triple Quad MS is calibrated with the Agilent ESI-L Low concentration Tuning mix. Source parameters: gas temperature 150°C, gas flow 10 L/min, nebulizer 45 psi, sheath gas temperature 325°C, sheath gas flow 12 L/min, capillary –2000 V, and delta EMV –200 V. Negative ion mode was used. Dynamic multiple reaction monitoring (dMRM) scan type is used with 0.07 min peak width, acquisition time is 24 min. Delta retention time of plus and minus 1 min, fragmentor of 40 eV, and cell accelerator of 5 eV are incorporated in the method. Data were pre-processed with Agilent MassHunter Workstation Quantitative Analysis for QQQ version 10.1, build 10.1.733.0.

#### Co-immunoprecipitation

HEK cells were transfected 24 h after plating. Mixed homemade PEI 3µl with 1µg of plasmid in 200µls of Opti MEM per well of a 6 well plate, incubated at room temperature for 20mins, then gently added to the cells. Cells were lysed (6 wells of a 6 well plate combined) per condition with RIPA buffer supplemented with protease inhibitor cocktail, sodium orthovanadate, MgCl, DNaseI and EDTA (final concentration of all these at 1mM). Lysates were incubated on ice for 20mins, then spun down at 15,000rpm for 15mins, with subsequent collection of the supernatant. 90% of supernatant volume were incubated with GFP beads (already washed according to manufacturer’s protocol) and the remaining 10% were used as input control. Incubation with GFP beads continued at 4°C for two hours while mixing. Afterwards, samples were washed three times according to manufacturer’s protocol and then bead-bound protein was diluted with 2X dye and boiled at 95°C for 10mins.

#### RNA extraction and RNA sequencing

Total RNA from proliferation assays was extracted using TRIzol (Invitrogen) using the manufacturer’protocol and the quality of the sample was analyzed using the 2100 bioanalyzer system (Agilent). Samples were then sequenced using the Element AVITI platform.

#### KPC Mice

*LSL-Kras^G12D/+^*;*LSL-Trp53^R172H/+^;Pdx1*-*Cre* (KPC) mice were bred by crossing the individual *LSL-Kras^G12D/+^*, *LSL-Trp53^R172H/+^*, and *Pdx1*-*Cre* strains. Triple mutant mice were palpated twice weekly for evidence of early tumors beginning at 8 weeks of age, followed by subsequent B-mode ultrasound screening using a VisualSonics 2100 Vevo High Resolution System. Following detection, tumors were monitored once weekly until reaching a mean diameter of 6 mm.

#### Oxylite Measurements

Intratumoral partial oxygen pressures in KPC mice (n=16) were measured using the OxyLite fluorescence quenching-based system (Oxford Optronics). Tumor-bearing KPC mice were anesthetized with 2% isoflurane in either air or pure oxygen. Hair was removed with depilatory cream around the abdomen and the tumor was visualized by ultrasound. A syringe with a 21G needle was attached to a stereotactic mount and inserted through the skin and abdominal wall. Real-time ultrasound imaging was used to visually guide the needle in-plane with the image through the center of the tumor until reaching the far edge. With the needle in place, the syringe was carefully removed and the bare-fiber oxygen-sensing OxyLite probe was then attached to the stereotactic mount and threaded through the needle bore until the probe tip was localized at the far edge of the tumor. The needle was fully retracted over the fiber and an initial pO_2_ measurement was taken at the far site. Prior to each measurement, the probe was allowed to equilibrate for 3-5 minutes until readings stabilized. After the initial reading, the fiber was retracted incrementally through the needle track, with readings taken every 1-2 mm, through the full depth of the tumor (ranged from 6-12 mm in diameter). Measurements within 1 mm of the edge of the tumor were excluded from the analysis since the needle frequently punctured the far wall of the tumor, allowing oxygen from the abdominal cavity into the wound (as made apparent by a sharp spike in readings). To compare tumor to normal tissue, partial pressures were also measured in pancreas and kidney of wildtype mice (n=4) kept under anesthesia using compressed air as vehicle. After completing measurements, mice were euthanized by isoflurane overdose and tissue was harvested for formalin fixation for 24 h prior to paraffin embedding. All tumors were verified as pancreatic ductal adenocarcinoma by a blinded observer experienced in mouse tumor pathology.

#### In vivo xenograft studies

In order to generate orthotopic xenograft tumors, survival surgeries were carried out and 1×10^5^ to 1000 tumor cells in 30–50 µl media/Matrigel mixtures (1:1) were implanted directly into the mouse pancreas using Panc1 and KP4 cell lines respectively. Body weights were measured and tumor growth was measured by high-resolution 3D ultrasound imaging weekly ^99^.

#### Single cell RNA sequencing

KPC samples used in Wasko et al. were submitted for Single-Cell RNA-Sequencing to the Sulzberger Genome Center. Single-cell sequencing data were processed using the Cell Ranger pipeline from 10X Genomics. FASTQ files were generated and aligned using the mouse transcriptome as a reference (v. gex-mm10-2020-A). ScRNA-seq profiles from each of the samples (both Controls and drug-treated) were quality controlled and filtered based on minimum and maximum UMIs per cell, (10^3^ and 10^5^, respectively) and the percentage of mitochondrial UMIs (max 25%). The resulting scRNA-Seq data were embedded in a Seurat object for normalization and scaling using the procedure outlined in ^100^. The optimal number of clusters was determined by the resolution-optimized Louvain algorithm, as described in ^101^. Unbiased inference of main cell types was performed using the SingleR package in combination with two mouse datasets as references contained in the celldex package (MouseRNAseqData ^102^ and ImmGenData ^103^). SingleR-inferred cell types were confirmed and refined after careful inspection of the most differentially expressed genes per cluster determined using a Wilcoxon Rank Sum test as implemented in the FindAllMarkers of the Seurat package. The malignancy status of the clusters of putative tumor cells was confirmed by performing inferCNV analysis ^104^. Sample-specific gene regulatory networks were inferred using the ARACNe3 algorithm ^105^ applied to the malignant cells of the Control samples. ARACNe3 networks were then integrated across samples, in order to create consensus regulatory models. The following ARACNe3 parameters were used: 100 subnetworks, 0.25 FDR. Drug response gene expression signatures were computed by comparing the expression level of each gene in each drug-treated sample with respect to the average of the same gene in all vehicle controls. The resulting signatures were then converted into protein activity using the NaRnEA algorithm ^105^, in combination with the Malignant Cells regulatory network. This produces a NES – a measure of the statistical significance – and a proportional enrichment score (PES) – a measure of effect size, for each inferred regulator.

---

**Supplementary Figure 1.**
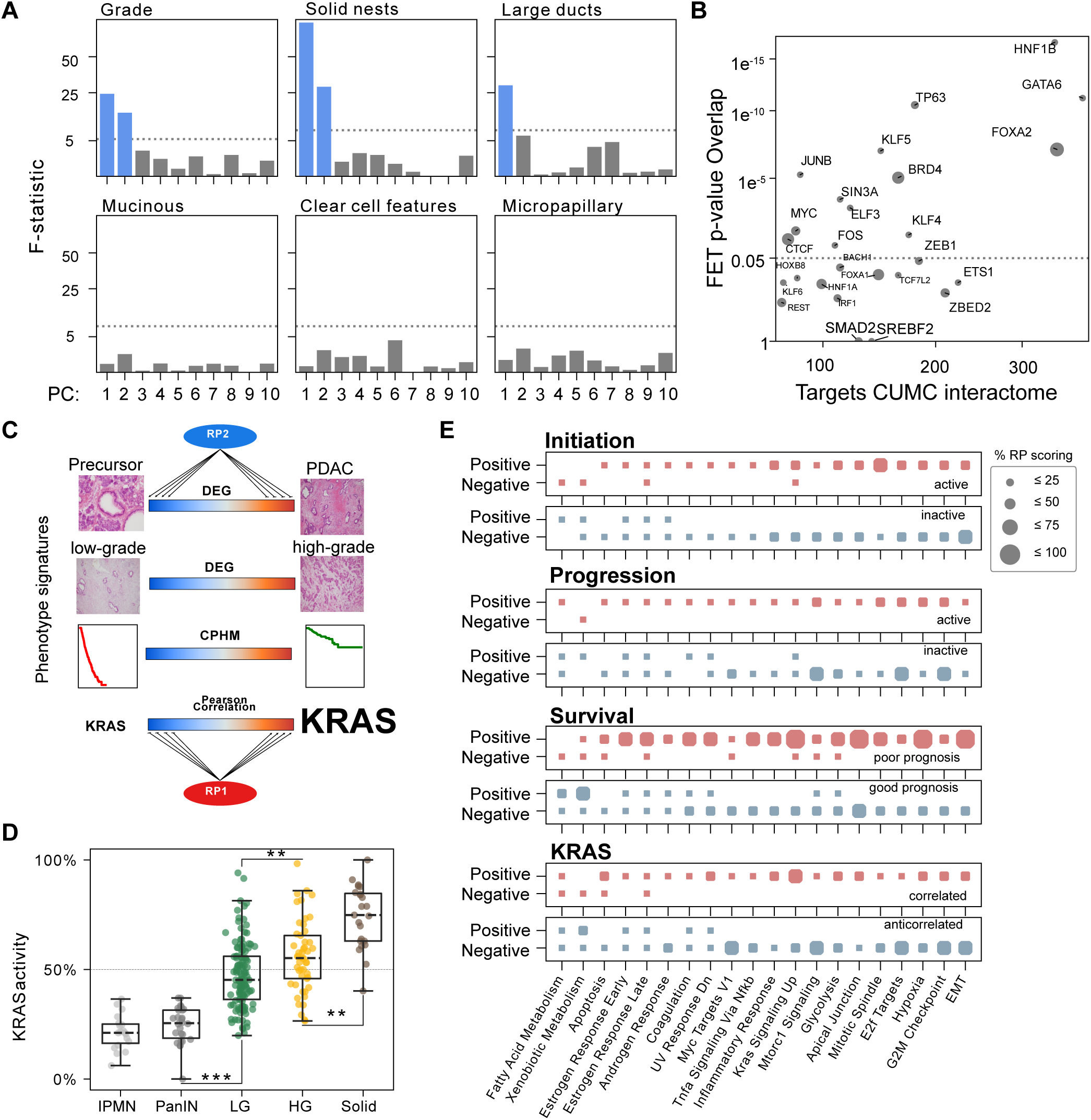
(**A**) F-statistics (y-axis) from a linear model with the indicated principal component (PC, x-axis) as dependent variable and the indicated histopathological characteristic as the independent variable. Blue bars indicate significant results. (**B**) Scatter plot illustrating the relationship between number of ARACNe-inferred, context-specific targets for select regulatory proteins (RP, x-axis) and the significance of their overlap with targets deduced from publically available ChIP experiments as assessed by a two-tailed Fisher’s Exact test. Circle size represents the number of publically available ChIP experiments (range = 1-4) (**C**) Schematic illustrating the assessment of regulatory protein activity in various genome-wide phenotype signatures. (**D**) KRAS protein activity across the indicated histological PDAC stages. Pairwise p-values are derived from a post hoc Dunn test after Kruskal-Wallis one-way analysis of variance. LG = Low grade; HG = High grade. (**E**) For each of 4 phenotypic signatures, regulons from the top 20 RP up (red color) and down (blue color) were divided into positive and negative targets (y-axis) which were then scored for overlap with the indicated HALLMARK gene sets (x-axis). Circle size represents the fraction of RPs (20 each) whose positive and negative regulon tail, respectively, exhibits significant overlap as assessed by a two-tailed Fisher’s Exact test. **: p ≤ 0.01, ***: p ≤ 0.001 In boxplots, the box ranges from Q1 (the first quartile) to Q3 (the third quartile) of the distribution and the range represents the IQR (interquartile range). The median is indicated by a dashed line across the box. The “whiskers” on box plots extend from Q1 and Q3 to 1.5 times the IQR.

**Supplementary Figure 2.**
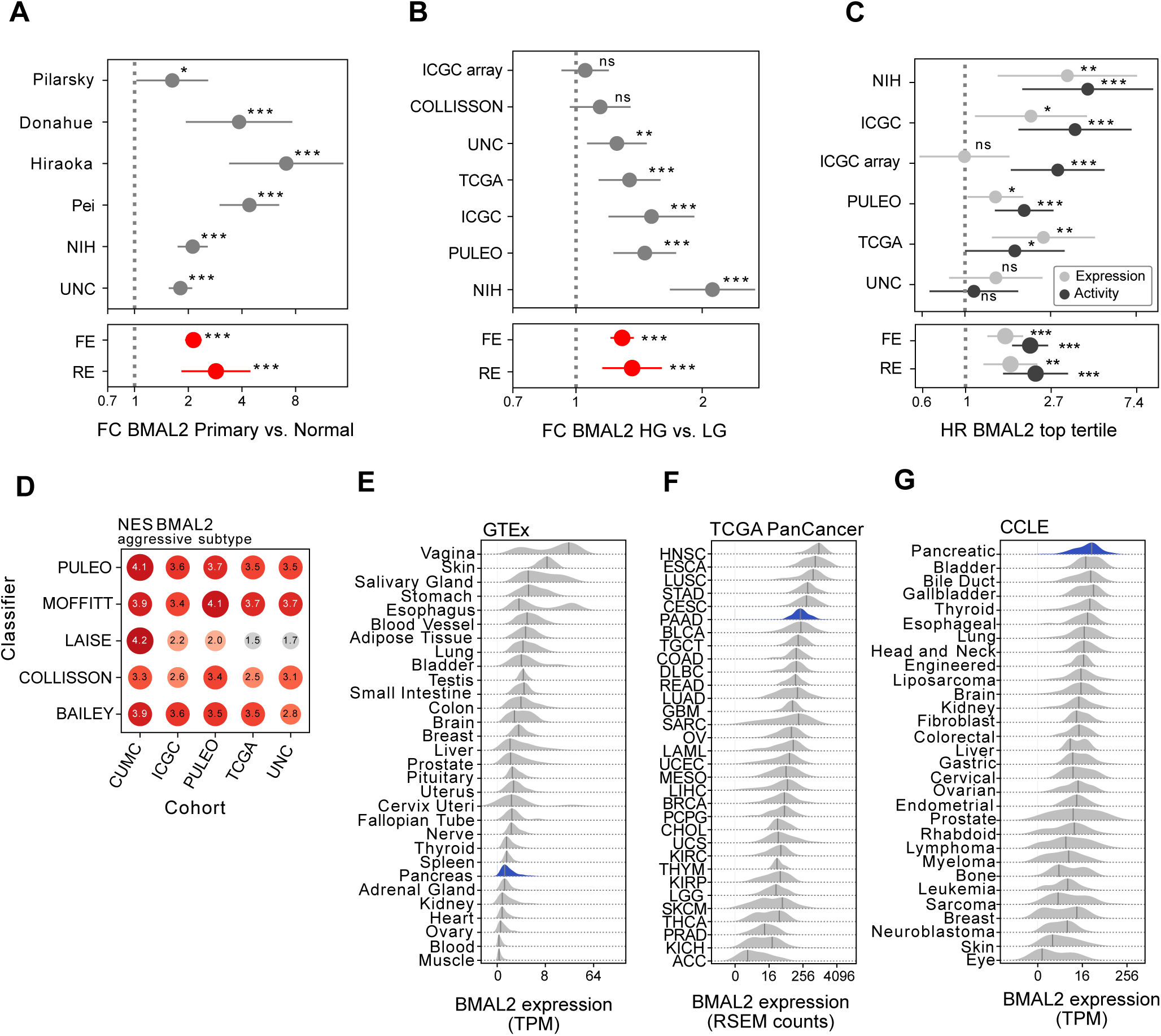
Fold changes and their 95% confidence interval between (**A**) primary tumors and adjacent normal tissue, and (**B**) high-grade and low-grade tumors for the indicated data sets (y-axis). (**C**) Hazard ratios and their 95% confidence interval for patients belonging to the highest BMAL2 activity and expression tertile, respectively. Lower panels in (**A-C**) summarize the meta-analytic estimate from a fixed (FE) and random effects (RE) model, respectively. (**D**) BMAL2 activity (Normalized Enrichment Score, NES) in the most aggressive vs. least aggressive subtype as determined by the indicated classification scheme (y-axis) in each of the indicated data sets (x-axis). BMAL2 expression in the indicated units across (**E**) various normal tissues, (**F**) 33 primary tumor cohorts profiled by the TCGA Pan-Cancer project, and (**G**) cancer cell lines derived from various lineages profiled by the CCLE project. *: p ≤ 0.05 **: p ≤ 0.01, ***: p ≤ 0.001, ns: not significant

**Supplementary Figure 3.**
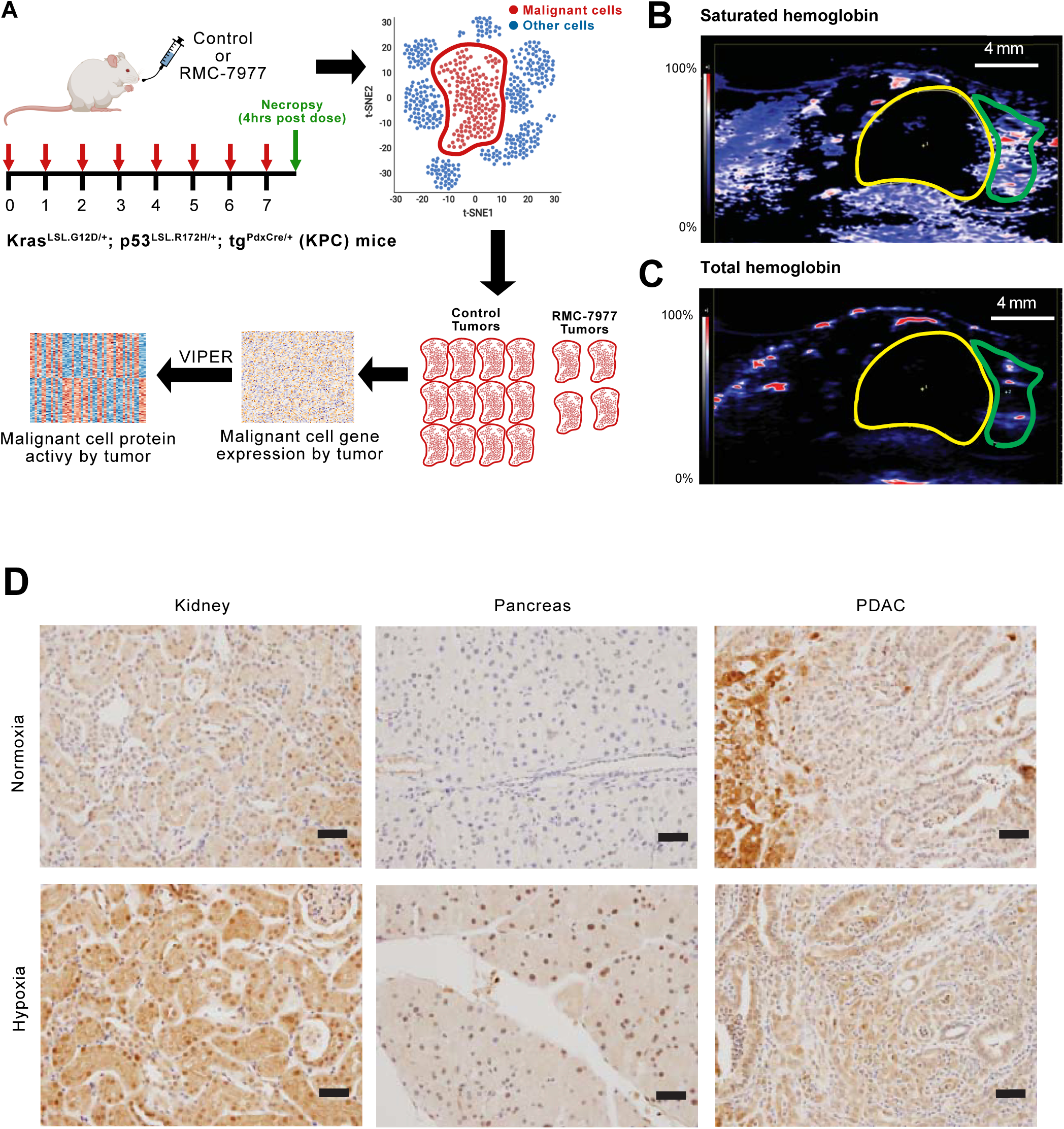
(A) Experimental design of the single-cell RNA sequencing experiment experimental setup. KrasLSL.G12D/+; p53LSL.R172H/+;Pdx1-Cretg/+(KPC) mice were treated with either vehicle or RMC-7977 (50 mg/kg q.2d. p.o.) for 7 days. Tissue was collected and processed for scRNA sequencing. Finally, tumor cells were analyzed and inferred differential activity of proteins was calculated based on the expression of their downstream target genes. (B) Heatmap of photoacoustic data from a representative KPC tumor, where red indicates high % blood oxygenation and blue represents low % blood oxygenation. Based on anatomical data from a co-registered B-mode image (not shown), tumor is outlined in yellow and adjacent normal pancreas is outlined in green. (D) Heatmap of total hemoglobin content from a representative KPC tumor (yellow) and adjacent normal pancreas (green). (D) Anti-pimonidazole (hydroxprobe) IHC stainings of FFPE blocks of normal kidney (left), normal pancreas (middle) and KPC tumors (right) are shown from mice under normoxic (top) or hypoxic (bottom) conditions. Scale bars = 20um.

**Supplementary Figure 4.**
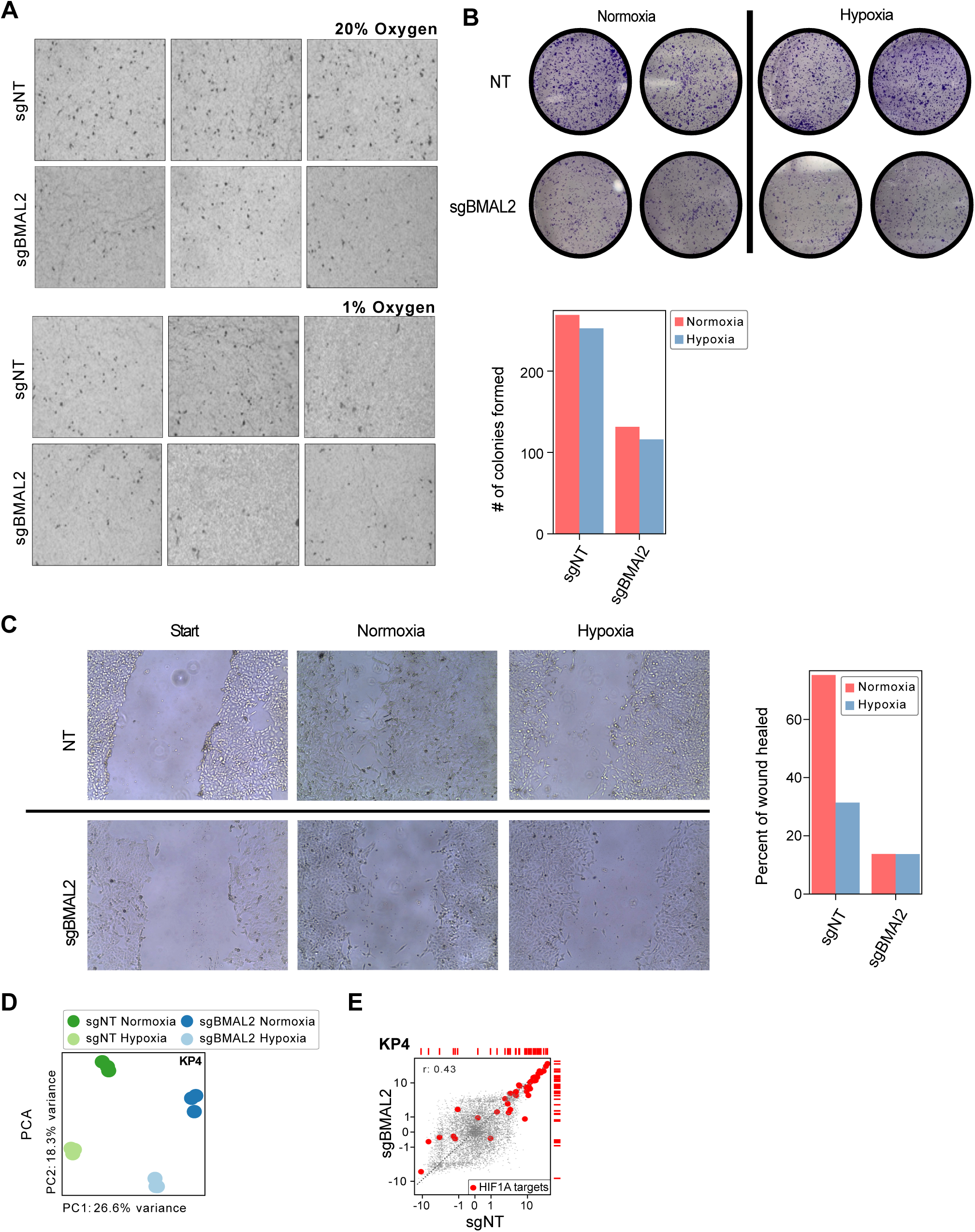
(**A**) Representative transwell migration assay pictures of PANC1 cells carrying sgNT or sgBMAL2 under normoxic or hypoxic conditions. (**B**) Cell clone formation assessed using a plate-based assay in KP4 cells carrying sgNT or sgBMAL2 under normoxic or hypoxic conditions (top panel) with the quantification (bottom panel) (**C**) Effect of BMAL2 knock out on cell migration was detected using a scratch assay. The scratching area was photographed at starting point and 24h under normoxia or hypoxia conditions (left panel) and quantified (right panel). (**D**) Principal component analysis (PCA) based on gene expression of the indicated cell line carrying sgNT or sgBMAL2 in hypoxic or normoxic environments, respectively. (**E**) Scatter plot illustrating the relationship of a genome-wide transcriptional hypoxia signature found in the indicated cell lines carrying sgNT (x-axis) and sgBMAL2 (y-axis) sgRNA, respectively. Red circles mark a set of HIF1A reporter genes described previously (Ref. 57)

